# Allosteric regulation of Thioesterase Superfamily Member 1 by free fatty acids and lysophosphatidylcholine

**DOI:** 10.1101/2020.02.18.954917

**Authors:** Matthew C. Tillman, Norihiro Imai, Yue Li, Manoj Khadka, C. Denise Okafor, Puneet Juneja, Akshitha Adhiyaman, Susan J. Hagen, David E. Cohen, Eric A. Ortlund

## Abstract

Non-shivering thermogenesis occurs in brown adipose tissue to generate heat in response to cold temperatures. Thioesterase superfamily member 1 (Them1) is transcriptionally upregulated in brown adipose tissue upon cold exposure and suppresses thermogenesis to conserve energy reserves. Them1 hydrolyzes long-chain fatty acyl-CoAs, preventing their use as fuel for thermogenesis. Them1 contains a C-terminal StAR-related lipid transfer domain (StarD) with unknown ligand or function. By complementary biophysical approaches, we show that StarD binds to long-chain fatty acids, products of Them1’s enzymatic reaction, as well lysophosphatidylcholine (LPC), which activate thermogenesis in brown adipocytes. Certain fatty acids stabilize the StarD and allosterically enhance Them1 catalysis of acyl-CoA, whereas 18:1 LPC destabilizes and inhibits activity, which we verify in cell culture. Additionally, we demonstrate that the StarD functions to localize Them1 near lipid droplets. These findings define the role of the StarD as a lipid sensor that allosterically regulates Them1 activity and localization.

## Introduction

Brown adipose tissue (BAT) mediates non-shivering thermogenesis in both mice ^1^ and humans ^1, 2^. A key function of non-shivering thermogenesis is to maintain core body temperature upon exposure to cold ambient temperatures. Because high rates of caloric consumption are required to generate heat, pharmacologic approaches to increasing BAT mass and activity are viewed as promising objectives in the management of obesity and related metabolic disorders ^3^.

In addition to activating thermogenesis in BAT, cold ambient temperatures lead to the transcriptional upregulation of genes that regulate energy expenditure including Thioesterase superfamily member 1 (synonyms brown fat inducible thioesterase (BFIT), steroidogenic acute regulatory lipid transfer-related domain 14 (StarD14) and acyl-Coa thioesterase 11 (Acot11)) ^4, 5^. Expression is induced upon cold exposure, and this was originally believed to contribute to the thermogenic output of BAT ^5^. However, rather than promoting energy use, Them1 proved to suppresses thermogenesis, thereby reducing the energy output of mice ^6^. Mechanistically, Them1 hydrolyzes long-chain fatty acyl-CoAs that are derived from endogenous lipid droplets within brown adipocytes, preventing their use as fuel for thermogenesis ^7, 8^. The genetic ablation of Them1 enhances the thermogeneic output of mice and protects against diet-induced obesity and metabolic disorders ^6^.

Them1 comprises two N-terminal thioesterase domains that hydrolyze acyl-CoA and a C-terminal StarD. The Them1 StarD is a member of the larger StarD family which is characterized by a highly conserved ∼210 amino acid sequence found in both plant and animal proteins ^9, 10^.

Lipid ligands and functional roles have been proposed for several of the StarDs. For instance, cholesterol, 25-hydroxycholesterol, testosterone, phosphatidylcholine, phosphatidylethanolamine and ceramides bind to STARD1/STARD3/STARD4/STARD5 ^11, 12^, STARD5 ^13^, STARD6 ^14^, STARD2/STARD7/STARD10 ^15–17^, STARD10 ^17^ and STARD11 ^18^, respectively. StarD1 and StarD3 bind and transport cholesterol to the mitochondria for steroidogenesis ^11, 19, 20^. StarD2, StarD7, and StarD10 bind phosphatidylcholine and influence membrane lipid compositions ^11, 16, 17, 21^. In the case of Them1, it is not known whether the StarD binds to lipid or whether lipid recognition plays a role in regulating thermogenesis.

We have shown that the Them1 StarD is necessary for full catalytic activity of the acyl-CoA thioesterase domains in the hydrolysis of long-chain fatty acyl-CoAs; purified recombinant acyl-CoA thioesterase domains alone exhibited significantly attenuated catalytic activity, and this was restored upon the addition of purified recombinant StarD ^7^. This phenomenon is not restricted to Them1, evidenced by increased activity of the long-chain fatty acyl-CoA thioesterase, Them2, in the presence of StarD2 ^16, 22, 23^. This study examines the effect of the StarD in brown adipocytes and explores the mechanism by which the StarD regulates acyl-CoA thioesterase (Acot) activity. We identify long-chain fatty acids and lysophosphatidylcholines (LPCs) as ligands for the Them1 StarD. Certain fatty acids allosterically enhance, whereas 18:1 LPC inhibits Them1 activity in a StarD-dependent manner. We further verify that 18:1 LPC relieves suppression of fatty acid oxidation by Them1 in brown adipocytes in an immortalized brown adipose cell line, in keeping with allosteric inhibition of Them1. Additionally, we discover the StarD is necessary for localizing Them1 to the lipid droplet.

## Results

### Them1 StarD binds long-chain fatty acids

To identify possible ligands for the Them1 StarD, we used affinity purification coupled with mass spectrometry following exposure of recombinantly expressed 6xHis tagged Them1 StarD to mixed-lipid liposomes (Fig. 1A). Because fatty acids are generated as a product of Them1’s enzymatic reaction, we reasoned that these products may bind to the StarD to facilitate product release. Our initial strategy therefore involved a targeted quantitative free fatty acid assay ^24^. This identified 15 fatty acid species that copurified with the Them1 StarD (Fig. 1B).

**Figure 1.**
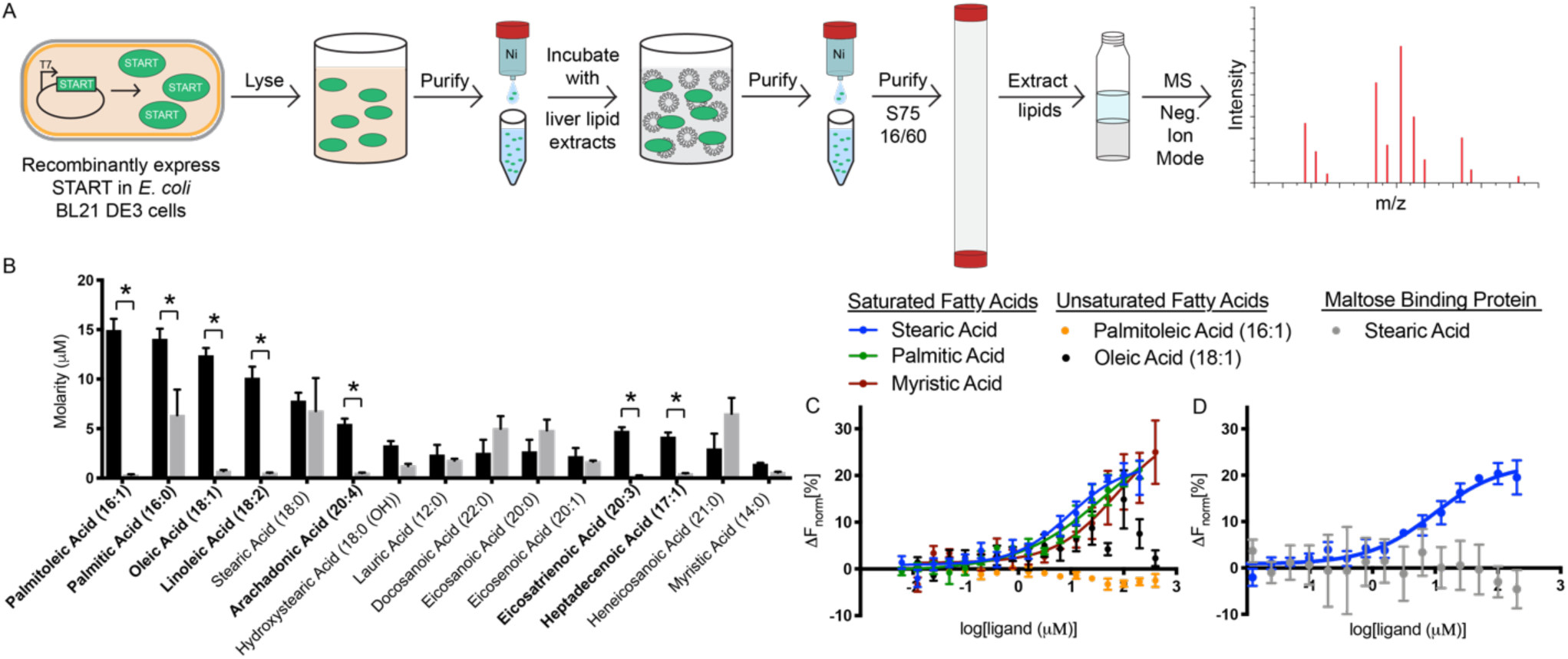
StarD of Them1 binds to long-chain fatty acids. *A.* Schematic of affinity purification mass spectrometry protocol. *B.* Concentration of 15 identified fatty acids bound to 1 milligram of Them1 StarD (black) or MBP (gray) as determined by negative ion mode mass spectrometry through normalization to deuterated fatty acid standard. Bars indicate average of three technical replicates. Error bars display standard error of mean. Statistical analyses were conducted using 2-way ANOVA with Sidak’s multiple comparisons test. *P<0.01. *C-D.* Fatty acid binding assay using microscale thermophoresis (MST). *C.* Them1 StarD labeled with Monolith RED-tris-NTA was kept constant at 50 nM, while the concentration of stearic acid (blue), palmitic acid (green), myristic acid (red), oleic acid (black), and palmitoleic acid (orange) were varied between 6.1 nM – 400 μM. Following an overnight incubation at 4 °C, StarD-FA solutions were loaded in standard Monolith NT.115 Capillaries (NanoTemper Technologies) and measured using a Monolith NT.115 instrument (NanoTemper Technologies). An MST-on time of 5 s was used for analysis, and baseline corrected normalized fluorescence values (ΔF_norm_[%]) were plotted against fatty acid concentration. Curves were fit with a nonlinear regression model and Kd’s are reported in Table 1 (n = 3 independent measurements, error bars represent the standard error of the mean). *D.* MST binding assay via titration of stearic acid (6.1 – 200 μM) into Monolith RED-tris-NTA labeled Them1 StarD (blue) or MBP (gray) held constant at 50 nM. Procedure and analysis were conducted the same as described previously.

**Table 1.** Affinity of fatty acids for Them1 START domain determined by MST.

| **Saturated Fatty Acids** |  |
| --- | --- |
| Myristic Acid (14:0) | 70.8 μM |
| Palmitic Acid (16:0) | 45.9 μM |
| Stearic Acid (18:0) | 9.5 μM |
| **Unsaturated Fatty Acids** |  |
| Palmitoleic Acid (16:1) | N.A. |
| Oleic Acid (18:1) | N.A. |

Seven of the identified fatty acids were significantly enriched within the StarD samples over our negative control (maltose binding protein), namely, palmitoleic acid (16:1), oleic acid (18:1), linoleic acid (18:2), palmitic acid (16:0), arachidonic acid (20:4), eicosatrienoic acid (20:3), and heptadecenoic acid (17:1) (Fig. 1B). The StarD showed a preference for unsaturated fatty acids with a tail length from 16 to 20 carbons, but also bound to saturated fatty acids, such as palmitic acid (16:0), and to a lesser degree myristic acid.

To determine the fatty acid binding affinity, we attempted to use traditional fatty acid binding assays that rely on competitive displacement of a fluorescent probe such as 1-anilinonaphthalene-8-sulfonic acid (1,8-ANS) ^25, 26^; however, the StarD did not bind to any of the probes tested. Therefore, we developed a fatty acid binding assay using microscale thermophoresis, which detects alterations in fluorescence along a temperature gradient induced by ligand binding to fluorophore-labeled protein ^27, 28^. We tested the top three fatty acids that copurified most abundantly with the StarD from lipid extracts, namely, palmitoleic acid (16:1), palmitic acid (16:0), and oleic acid (18:1). In contrast to expectations, only palmitic acid generated a binding curve (Fig. 1C). Since palmitic acid is a saturated fatty acid, we also tested myristic acid (14:0) and stearic acid (18:0); two saturated fatty acids that also copurified with the StarD in our MS analysis. Both generated binding curves (Fig. 1C). Binding for fatty acid species occurred in the µM range (Table 1). To ensure these alterations in thermophoresis were not due to micelle formation at high fatty acid concentrations, we titrated stearic acid (critical micellar concentration ≈ 300 μM ^29^) which is most prone to form micelles, into fluorescently labeled maltose binding protein and observed no changes in thermophoresis (Fig. 1D). These results suggest specific binding of saturated long-chain fatty acid to the Them1 StarD.

There were some discrepancies in the fatty acids identified to bind to the StarD by affinity purification-MS and by microscale thermophoresis techniques (Figs. 1C, D). This may be explained, at least in part, by differences in presentation of lipids to the StarD depending on the technique: the StarD was exposed to liposomes in our affinity purification-MS experiment whereas a single fatty acid was titrated into the StarD in the microscale thermophoresis experiment. Since unsaturated fatty acids copurified with the StarD and were detected by MS but did not bind in the microscale thermophoresis experiment, Them1 may have only accessed unsaturated fatty acids in the context of a membrane bilayer. Alternatively, unsaturated fatty acid binding may require the presence of other lipids to serve as intermediate ligands prior to lipid exchange.

### Fatty acids bind within the hydrophobic pocket of Them1’s StarD

To visualize the StarD-fatty acid complex, we attempted to generate crystals in the presence of a range of fatty acids, however, only myristic acid yielded crystals that diffracted. The myristic acid–StarD structure was solved in the P 1 21 1 space group to a resolution of 3.09 Å, with the asymmetric unit containing four StarD monomers (Fig. 2A). Refinement and model statistics are summarized in Table 2.

**Figure 2.**
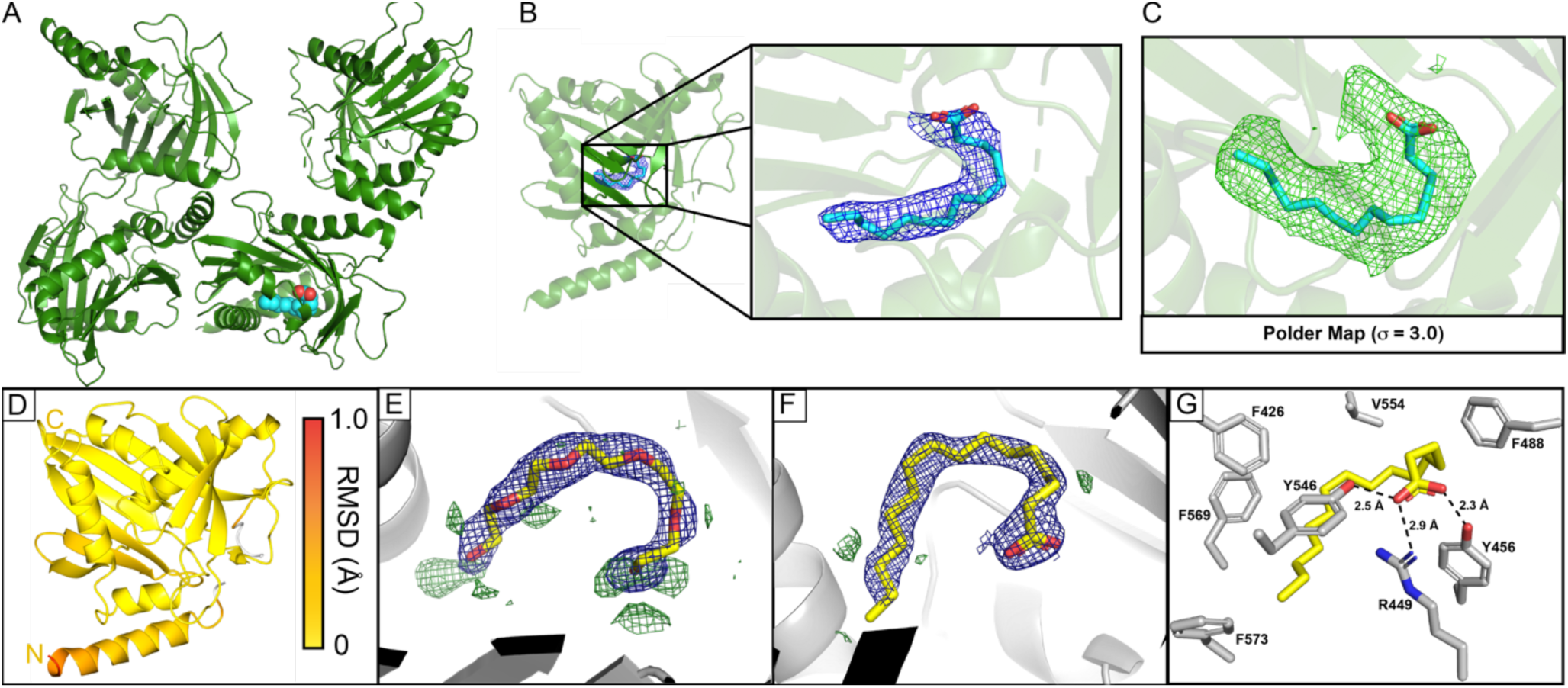
Fatty acids fit within crystal structure of Them1 StarD. *A.* Asymmetric unit cell of 3.09 Å structure of Them1 StarD contains 4 monomers (green), with one monomer bound to myristic acid (cyan). *B.* Zoomed in view of the lipid binding pocket of StarD where myristic acid (cyan) is modeled into 2F_o_-F_c_ map (blue mesh) contoured to *σ* = 1.0. *C.* Polder map (F_o_-F_c_ map with bulk solvent removed, green) contoured to *σ* = 3.0 of region surrounding fatty acid. Myristic acid (cyan) fits nicely within this density. *D.* ProSMART analysis conducted to determine r.m.s.d. between Cα backbone of myristic acid bound StarD and apo-StarD monomers. Root mean square deviations (range: 0–1.0 Å) between monomers were mapped onto myristic acid bound StarD structure with a color scale depicting low (yellow) to high (red) deviations. Unaligned regions are colored in white. *E.* PEG molecule (yellow) modeled into electron density of previous Them1 StarD crystal structure (PDB code: 3FO5). *F.* Palmitic acid (16:0) (yellow) modeled into refined density from previous structure of StarD. All 2F_o_-F_c_ maps displayed in blue and contoured to *σ* = 1.0. All F_o_-F_c_ maps displayed in green and contoured to *σ* = 2.0. *G.* Amino acids in close proximity surrounding palmitic acid. Distance between polar amino acids and carboxyl head group of fatty acid displayed as black dashed line

**Table 2.** Myristic Acid—Them1 START domain X-ray data collection and refinement statistics.

| **Data collection** | Myristic Acid—START |
| --- | --- |
| Space group | P 1 2<sub>1</sub> 1 |
| Cell dimensions |  |
| $a, b, c$ (Å) | 65.08, 70.98, 127.22 |
| $\alpha, \beta, \gamma$ (°) | 90, 96.13, 90 |
| Resolution (Å) | 42.17 – 3.09 (3.20 – 3.09) |
| CC<sub>1/2</sub> | 0.647 |
| R<sub>pim</sub> | 0.121 (0.71) |
| $I / \sigma I$ | 11.3 (1.4) |
| Completeness (%) | 98.97 (96.86) |
| Redundancy | 5.8 (5.3) |
| **Refinement** |  |
| Resolution (Å) | 3.09 |
| Unique reflections | 21201 (2040) |
| $R_{\text{work}} / R_{\text{free}}$ (%) | 21.76/27.72 (29.53/35.95) |
| No. non-hydrogen atoms | 7515 |
| Protein | 7490 |
| Ligands | 16 |
| Water | 9 |
| B-factors | 88.73 |
| Protein | 88.73 |
| Ligand | 101.73 |
| Water | 60.46 |
| Clashscore | 5.72 |
| R.m.s. deviations |  |
| Bond lengths (Å) | 0.004 |
| Bond angles (°) | 0.67 |
| Ramachandran favored (%) | 95.45 |
| Ramachandran allowed (%) | 4.32 |
| Ramachandran outliers (%) | 0.22 |
| PDB accession code | 6VVQ |

Values in parenthesis indicate highest resolution shell

One subunit contained continuous electron density within the interior of the domain that fit myristic acid (Fig. 2B). A polder map displayed clear density at a sigma level of 3.0, strongly supporting the presence of myristic acid binding at this location (Fig. 2C) ^30^. The other three subunits did not contain this continuous electron density; therefore, we suspect either no FA was bound, or that FA was bound with low occupancy or high conformational mobility. In this connection, myristic acid exhibited a higher B factor than the average B factor for the protein, suggesting the ligand had some mobility in the pocket (Table 2). Because we crystallized two different states of the StarD (with and without myristic acid) in one crystal, we compared the structures of these two states using ProSMART analysis ^31^. The root mean squared deviation (RMSD) between the apo monomers and the myristic acid bound monomer were mapped onto the structure of the StarD complexed with fatty acid, (one comparison in Fig. 2D; other comparisons in Supplemental Fig. 1A). There were no major conformational differences between the structures suggesting fatty acid does not induce an appreciable conformational change, albeit our data do not discern whether the apo domains are truly devoid of fatty acid. There were some dissimilarities in the N-terminal α-helix, but these changes are likely an artifact of crystal packing.

We next compared our structure with a structure of the Them1 StarD that was solved at higher resolution and in a different space group ^32^. Our structure exhibits the same overall conformation with an average RMSD of 0.4 Å over aligned atoms ^32^. The prior structure contained long, tubular electron density in the same location where we modeled a myristic acid. A buffer component (PEG molecule) was modeled into this density because no ligand was identified in the crystal structure (Fig. 2E) ^32^. We observed branched electron density characteristic of a fatty acid carboxyl-head group; therefore, we modeled palmitic acid, a highly abundant *E. coli* fatty acid that copurifies with the StarD, into this density (Fig. 2F) ^33^. Palmitic acid fit well within the density, providing strong support for placement of this fortuitously co-purified ligand. The palmitic acid carboxyl head group is contacted by polar residues arginine 449, tyrosine 456, and tyrosine 546 (Fig. 2G). The curved fatty acyl chain is enclosed by bulky, nonpolar amino acids, including phenylalanine 426, phenylalanine 488, valine 554, phenylalanine 569, and phenylalanine 573, which fully protect it from solvent (Fig. 2G).

To explain Them1’s unexpected preference for fatty acids, which are smaller-sized lipids than typically bind StarDs, we analyzed the lipid binding pockets of all StarDs of known structure. These share a long, continuous C-terminal α-helix that packs across the mouth of a U-shaped incomplete *β*-barrel, forming the empty interior (Supplemental Fig. 1C). The conformation of this helix is radically different in Them1, whereby a kink, enabled by a highly conserved glycine 564 and a steric clash from α-helix *α*0 (connecting thioesterase domain and the StarD), constricts the lipid-binding pocket ^32^. Supplemental Table 1 displays the surface area (Å^2^) and volume (Å^3^) of each lipid binding pocket, which are also shown graphically (Supplemental Fig. 1B). Them1 possesses a smaller interior cavity than STARD2 and STARD11, which bind to phosphatidylcholine (PC) and ceramide respectively. All other StarD proteins contain an interior pocket with similar area and volume, though a different shape, when compared with Them1 StarD. These other StarD proteins have resisted efforts at co-crystallization with their cholesterol and sterol-like ligands ^32, 34, 35^. Thus, the calculated size of the pocket size may not accurately reflect the ligand-bound state.

We superposed Them1 StarD with STARD2 bound to palmitoyl-linoleoyl phosphatidylcholine (PDB code: 1LN3) ^36^ and STARD11 bound to C16-ceramide (PDB code: 2E3P) ^36, 37^. In both instances, the interior cavity of the Them1 StarD was unable to accommodate the same ligands (Supplemental Fig. 1D). Them1 StarD contains some equivalent structural features that enable StarD2 and StarD11 to bind to their respective ligands, such as arginine 449 (StarD2 R78, StarD11 R442), which electrostatically interacts with the phosphate of phospholipids in StarD2 ^36^ and a water mediated hydrogen bond with a ceramide hydroxyl in StarD11 ^36, 37^. Additionally, these StarDs contain an acidic residue (Them1 D453, StarD2 D82, StarD11 E446) that participates in a salt bridge with the conserved arginine and engages in hydrogen bonding with the amide-nitrogen and hydroxyl of ceramide in StarD11 ^37^. However, Them1 lacks the aromatic cage found in STARD2 that consists of tryptophan 101, tyrosine 114, and tyrosine 155, which together engage in cation-*π* interactions with the quaternary amine of choline ^36^. Them1 only contains one structurally analogous aromatic residue (F488), though it also contains tyrosine 456 that could rotate and potentially occupy the same space as tryptophan 101 in STARD2 (Supplemental Fig. 1E). Additionally, Them1 does not conserve residues found in StarD11 (Y482, Q467, and N504) that participate in hydrogen bonding with ceramide ^37^.

Them1 lacks some necessary residues for recognition of the larger lipids present in StarD2 and StarD11, while coopting residues such as R449 and D453 to enable fatty acid binding.

### Them1 StarD binds to lysophosphatidylcholine

To test whether long-chain fatty acids were the only ligands for the StarD of Them1, we repeated our affinity purification mass spectrometry experiment using a shotgun MS approach that examined all major phospholipid classes. Several LPC species including 18:2, 18:1, 20:4, 20:3, and 22:4 were highly enriched in our StarD samples, but absent in our negative control samples (Fig. 3A-C). Them1 StarD samples were not enriched for any phosphatidylethanolamine, phosphatidylserine, or phosphatidylinositol species. Some sphingomyelin and phosphatidylcholine species were enriched in the StarD samples at low levels.

**Figure 3.**
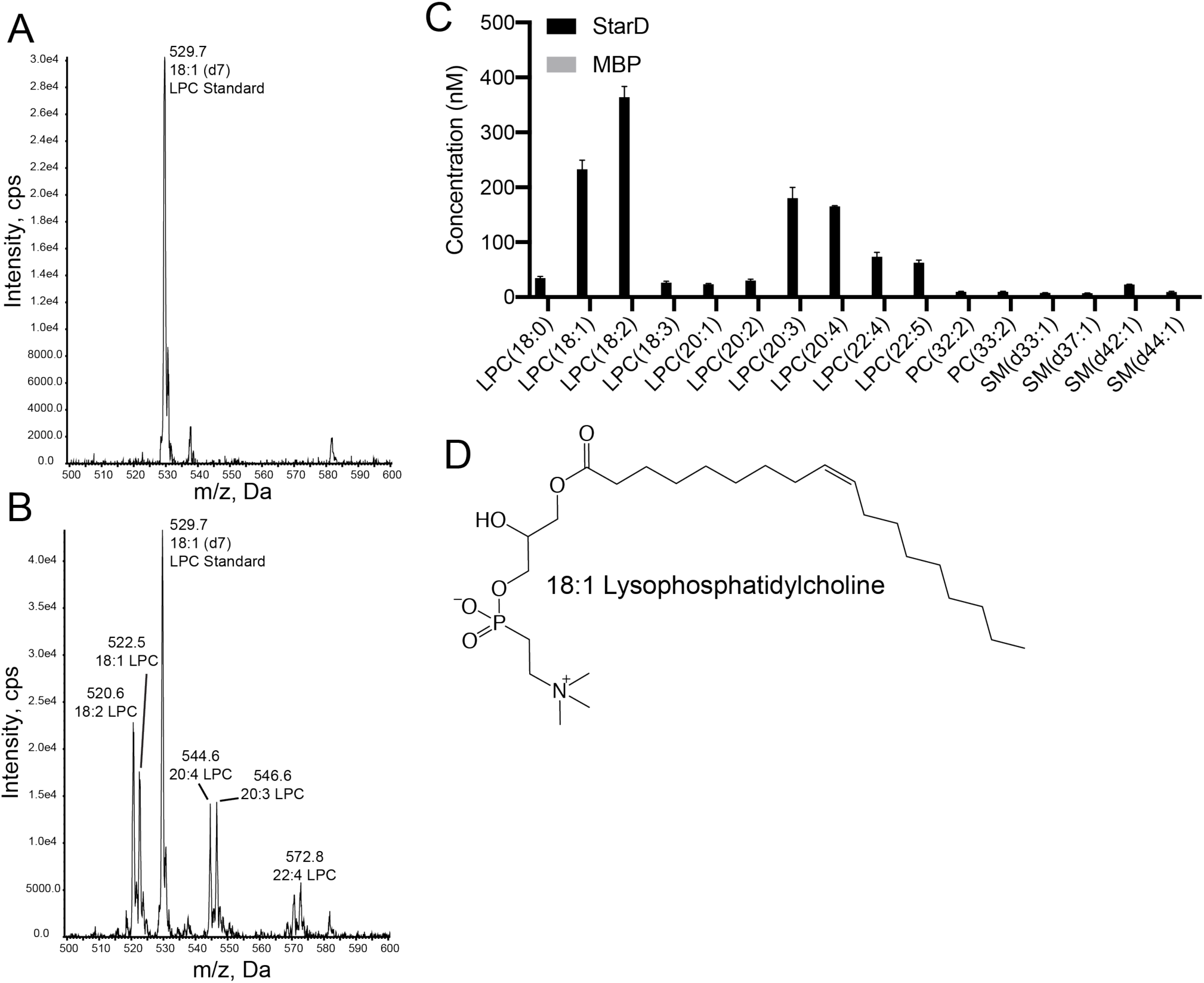
Them1 StarD domain binds to lysophosphatidylcholine. *A-B.* Mass spectra (precursor ion scan of *m/*z 184) of lipids containing a phosphatidylcholine head group that copurified with maltose binding protein (MBP) *(A)* and Them1 StarD *(B)*. Lysophosphatidylcholine 18:1 (d7) standard was added to each sample for quantification of lipid concentrations. *C.* Graphical analysis of identified lysophosphotidylcholine species. Bars are an average of three technical replicates. Lysophosphotidylcholine species were not detected in MBP samples. *D.* Chemical structure of 18:1 lysophosphotidylcholine.

LPC contains a single fatty acyl chain typically esterified at the *sn-1* position rather than the two fatty acyl chains present in phospholipids (Fig. 3D). It is an important signaling molecule that is implicated in the pathogenesis of cardiovascular disease, atherosclerosis, diabetes, and neurodegenerative diseases ^38^. LPC also functions in lipid droplet formation because it is a precursor, along with fatty acyl-CoAs, for phosphatidylcholine molecules that are required to expand the membrane monolayer that coats lipid droplet membranes ^39^. Upon stimulation of thermogenesis, levels of saturated LPC in brown adipocytes levels dramatically increase in brown adipocytes, which interestingly enhances thermogenesis ^40^.

### Fatty acids enhance while 18:1 LPC inhibits Them1 acyl-CoA thioesterase activity

Since the StarD was previously shown to alter the enzymatic activity of Them1 ^7^, we reasoned fatty acids and LPC species may regulate Acot activity through interaction with the StarD. To test this, we monitored myristoyl-CoA hydrolysis in the presence of fatty acid or 18:1 LPC. Incubation with either 25 μM myristic acid or palmitic acid enhanced the maximum enzymatic velocity of ΔNterm-Them1 lacking the intrinsically disordered N-terminus (Fig. 4A, red and green). This enhancement in activity is dependent upon the StarD (Fig. 4B). Incubation with stearic acid did not alter the activity of ΔNterm-Them1; however, stearic acid suppressed enzymatic activity of Them1 when the StarD was absent, indicating the StarD relieves the inhibitory effects of stearic acid (Fig. 4A-B, blue).

**Figure 4.**
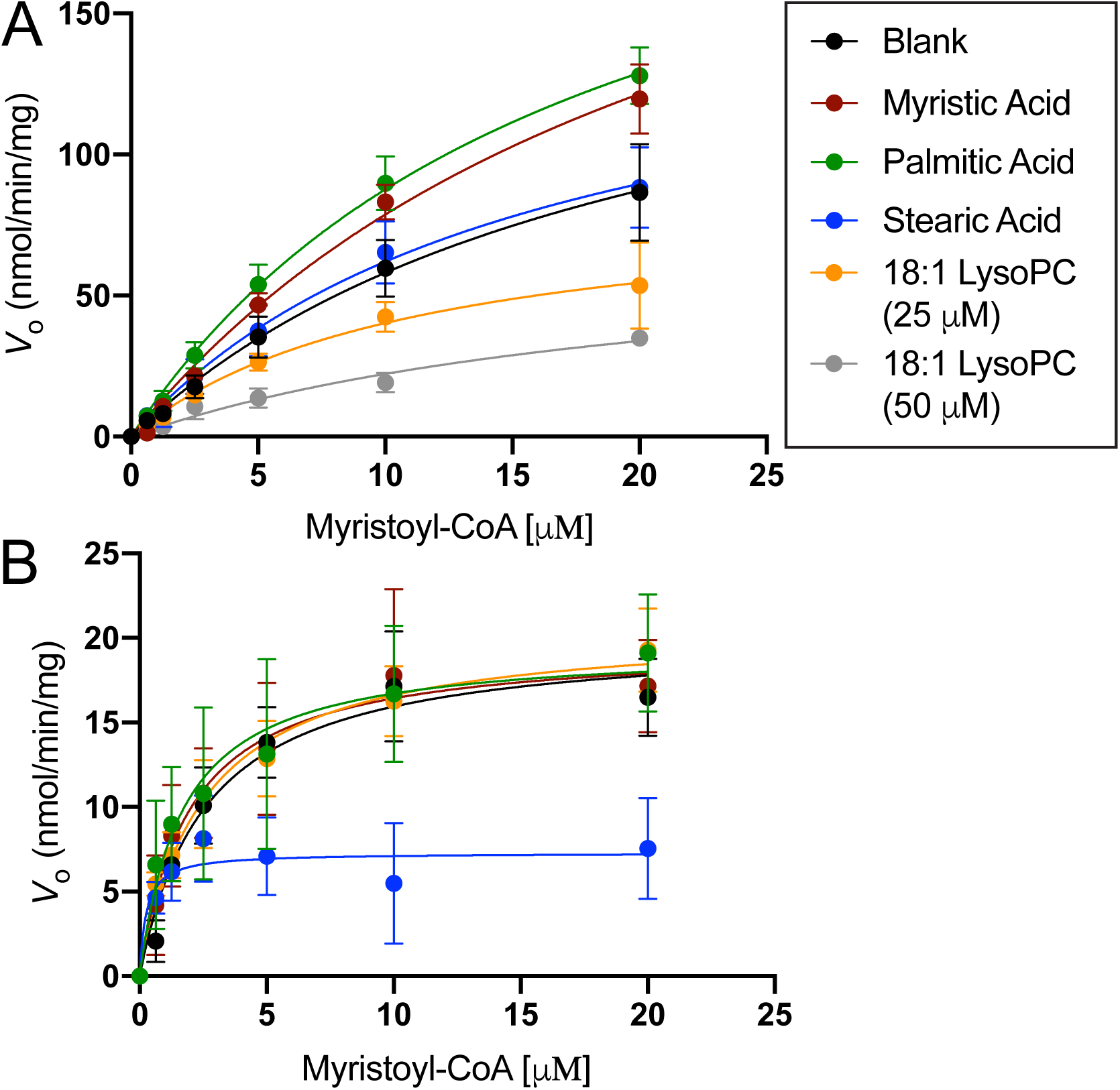
Fatty acids enhance while lysophosphatidylcholine inhibits Them1 activity in a StarD-dependent manner. ΔNterm-Them1 *(A)* and Δ Nterm-Them1_ΔStarD *(B)* (1 μM) were incubated with buffer (black) or 25 μM stearic acid (blue), palmitic acid (green), myristic acid (red), 18:1 lysophosphatidylcholine (orange), and 50 μM 18:1 lysophosphatidylcholine (gray) for 30 minutes at 37 °C prior to the addition of myristoyl-CoA. Saturation curves of *V_0_* plotted against increasing myrstoyl-CoA with solid lines indicating nonlinear analysis of the data. Each point corresponds to the average of a minimum of three replicates. Error bars represent standard error of the mean.

Surprisingly, incubation with 25 μM 18:1 LPC greatly inhibited ΔNterm-Them1 activity (Fig. 4A, orange) in a StarD dependent manner (Fig. 4B). To determine whether this inhibition was dose dependent, we incubated ΔNterm-Them1 with twice the concentration of 18:1 LPC (50 μM) and observed greater inhibition (Fig. 4A, gray).

### Them1 forms homotrimer containing a thioesterase domain core flanked by mobile StarDs

To understand how the StarD interacts with the thioesterase domains to influence catalytic activity we generated a model of full-length Them1 by joining the StarD structure with a homology model of the Them1 thioesterase domains created using the SWISS MODEL server ^41–45^ based on the structure of the ACOT12 thioesterase domains (56 % sequence identity and 69 % sequence similarity) ^46^. Both structures contained a common α-helix that resides at the C-terminus of the thioesterase domains model and N-terminus of the StarD structure. We aligned this overlapping α-helix to generate a full-length model of Them1 with similarities to a previously reported model of intact ACOT12 (Fig. 5A) ^46^.

We used single particle negative stain electron microscopy to obtain a low-resolution map of Them1 to fit our structural model. ΔNterm-Them1 was purified as a stable trimer as determined by size exclusion chromatography and analytical ultracentrifugation (Supplemental Fig. 2A-B). Negative stain electron microscopy revealed a homogenous distribution of ΔNterm-Them1 trigonal particles with a diameter of 17 nm spread across the grid (Supplemental Fig. 2C). We generated 2D-classifications using Relion 3.0 ^47^ that revealed a trimeric complex consisting of a large spherical body flanked by three protruding loves representing 3 fold symmetry (Fig. 5B, all class averages in Supplemental Fig. 2D). A 3D initial model was generated and refined using particles from selected class averages that revealed the trimeric complex (Fig. 5C). The central density of the 3D reconstruction accommodates the core heterotrimeric thioesterase domains; however, fitting the StarD within the peripheral density required relaxing the linker geometry of our model and independently fitting this domain (Fig. 5C-D).

The negative stain 2D class averages also revealed conformational flexibility between the StarD and thioesterase domains. The main central body of the map that accommodates the thioesterase domains is identical in all 2D class averages; however, the three exterior lobes that accommodate the StarDs are not perfectly arranged in a threefold symmetric frame in a few of the 2D class averages as seen in reprojections using C3 symmetry (Supplemental Fig. 2E). We postulate this flexibility is the result of a less ordered region lying between the domains (residues 365 – 383, *H. sapiens*), which was also disordered in our crystal structure of the StarD.

**Figure 5.**
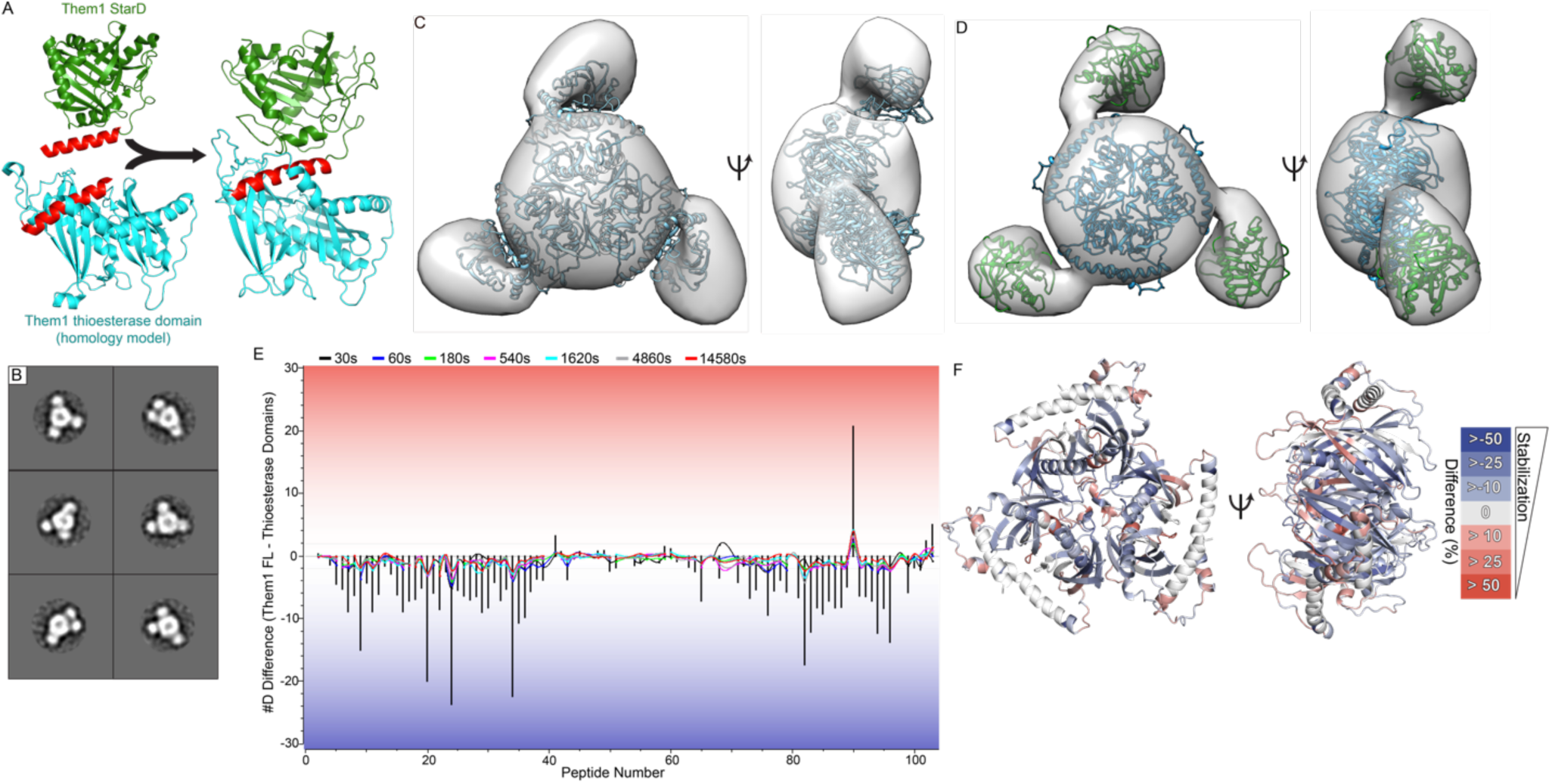
Them1 forms homotrimer with thioesterase domain core and flanking StarDs. *A.* Generation of full-length Them1 model through structural alignment of linker helix (red) present in crystal structure of Them1 StarD (green) and homology model of Them1 thioesterase domains (blue). *B.* Subset of 2D class averages of Them1 using negative stain electron microscopy generated by Relion 3.0. *C.* 3D reconstruction of Them1 derived from subset of 2D class averages. Trimeric Them1 model fit into 3D reconstruction using Chimera. *D.* Separate StarDs (green) and thioesterase domains (blue) modeld into 3D reconstruction using Chimera. *E.* Butterfly plot overlayed with a residual plot displaying difference in deuterium uptake between ΔNterm-Them1 and ΔNterm-Them1_ΔStarD. Colored lines depict deuterium uptake difference (y-axis) for peptides (x-axis) at each time point (black: 30s, blue: 60s, green: 180s, pink: 540s, gray: 1620s, cyan: 4860s, red: 14580s). Bars display summed difference in deuterium uptake over all time points for each peptide. Negative values (blue) mean that removal of StarD increases the incorporation of deuterium for the cooresponding peptide. *F.* Percentage difference in deuterium uptake (Them1 – Them1_ΔStarD) at 60 seconds mapped onto homology model of Them1 thioesterase domains. Negative values (blue) indicate less deuterium exchange (greater protection) in intact Them1, indicating that StarD stabilizes the thioesterase domains.

### Them1 StarD stabilizes the thioesterase domains

To determine how the thioesterase and StarDs interact, we performed hydrogen-deuterium exchange MS (HDX-MS). This technique identifies regions of flexibility and rigidity by measuring the rate of exchange of deuterium with amide protons; high deuterium uptake signifies areas of flexibility and high solvent exposure, whereas low deuterium uptake signifies areas of rigidity and low solvent exposure ^48^. Comparison of deuterium uptake of ΔNterm-Them1 with ΔNterm-Them1-ΔStarD reveals a dramatic stabilizing effect driven by the StarD, as evidenced by a great reduction of deuterium incorporation throughout the thioesterase domains in the presence of the StarD (Fig. 5E-F). Heat maps showing identified peptides for each construct are provided in Supplemental Fig. 3. Although there is flexibility between the thioesterase and StarDs, the StarD significantly stabilizes the thioesterase domains.

### Fatty acids stabilize while 18:1 LPC destabilizes StarD

Incubation with myristic acid, palmitic acid, or stearic acid did not alter the thermal melting temperature (*T_m_*) of the StarD, as monitored by differential scanning fluorimetry (DSF); however, this is relative to StarD that copurifies with fatty acids from *E. coli* (Fig. 6A).

**Figure 6.**
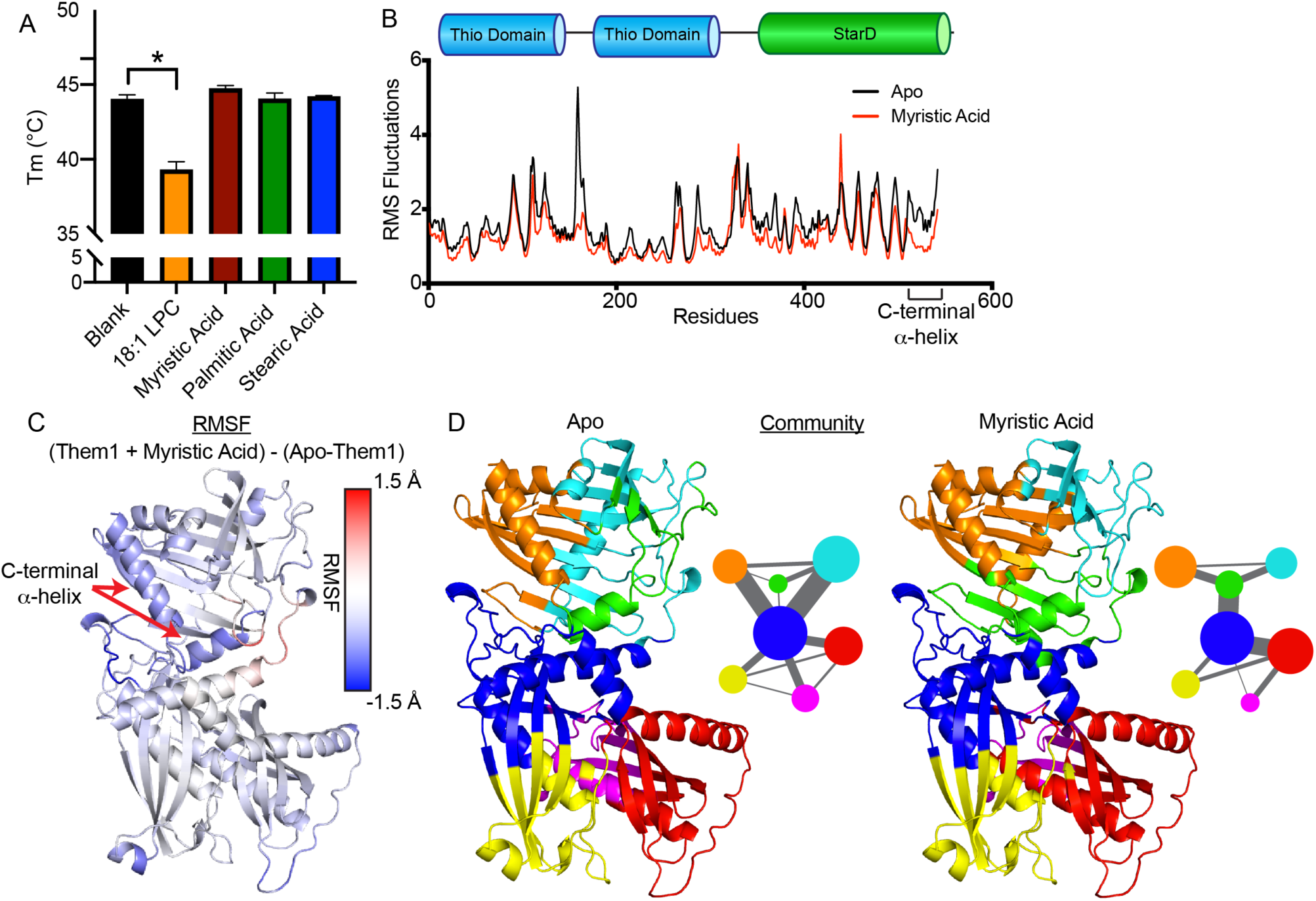
Fatty acids stabilize while 18:1 LPC destabilizes the StarD. *A.* Differential scanning fluorimetry of StarD incubated with either buffer (black), 18:1 LPC (orange), myristic acid (red), palmitic acid (green), and stearic acid (blue). LPC 18:1 significantly destabilizes the StarD. Bars depict average of three replicates. Error bars denote standard error of the mean. One-way ANOVA and Dunnett’s multiple comparisons test were used to analyze StarD data. *B-D.* Molecular dynamics simulation for apo-Them1 and myristic acid bound Them1 over 500 ns. *B.* Root mean squared fluctuations (RMSFs) across Them1 residues for the apo (black) and myristic acid bound (red) states. *C.* Color coordinated difference in RMSFs between myristic acid bound and apo states (Them1-MYR – Them1-Apo) mapped onto full-length model of Them1. Myristic acid stabilizes the C-terminal α-helix; blue color corresponds to lower RMSFs in myrstic bound state than apo state. *D.* Community analysis that identifies residues that move in coordinated fashion thorughout the simulation. Circle size depicts number of residues within community and width of lines corresponds with strength of communication between communities. Myristic acid alters size and connectedness of communities.

Strikingly, 18:1 LPC destabilized the StarD by nearly 5 °C, which is in-line with the reduced catalytic activity driven by this ligand (Fig. 6A).

To determine how fatty acids enhanced Them1 activity, we performed 500 ns molecular dynamics simulations on Them1 comparing apo vs lipid-bound Them1. The presence of myristic acid substantially lowered the root mean squared fluctuations in the StarD and in some parts of the thioesterase domains, suggesting myristic acid generated a more stable complex than apo-Them1 (Fig. 5B-C). The C-terminal α-helix of the StarD, which plays a role in StarDs binding lipids ^49, 50^, was significantly stabilized by myristic acid (Fig. 5B-C). Next, we performed a community analysis, which identifies groups of residues that move in a coordinated manner throughout the simulation. Myristic acid significantly altered the communities within the StarD, changing their size and connectedness (Fig. 5D). In the apo state, multiple communities interface with the blue community of the thioesterase domains, but myristic acid shifts the communities so that only one is connected to the thioesterase domains (Fig. 5D). Although there are fewer connections between the communities in the myristic bound state, there are stronger connections linking the communities in the StarD (green) with communities within the thioesterase domains where acyl-CoA hydrolysis occurs (blue-red), potentially yielding a more active enzyme (Fig. 5D). Taken together, these data suggest fatty acids allosterically enhance Acot activity through stabilizing the StarD and altering dynamics within the thioesterase domains, while 18:1 LPC inhibits Acot activity through destabilizing the StarD.

### 18:1 LPC reverses Them1-mediated suppression of fatty acid oxidation

To test the effect of the StarD on Them1’s capacity to suppress thermogenesis in live cells, we measured the oxygen consumption rate (OCR) of mouse-derived immortalized brown adipocytes (iBAs) that were transduced with EGFP alone, EGFP-tagged full-length Them1 (FL-Them1-EGFP), or a EGFP-tagged truncated variant containing only the N-terminal thioesterase domains (Them1_ΔStarD-EGFP) (Fig. 7A-B). We induced thermogenesis in the iBAs using norepinephrine (NE), which increased the OCR over basal levels (Fig 7B, green). As expected, Them1 suppressed NE-induced respiration (Fig. 7B, black). However, removal of the StarD did not reduce Them1-mediated suppression of OCR values, but slightly enhanced Them1 activity (Fig. 7B, red). The StarD in part attenuated Them1 activity, suggesting a role for feedback regulation by the StarD.

**Figure 7.**
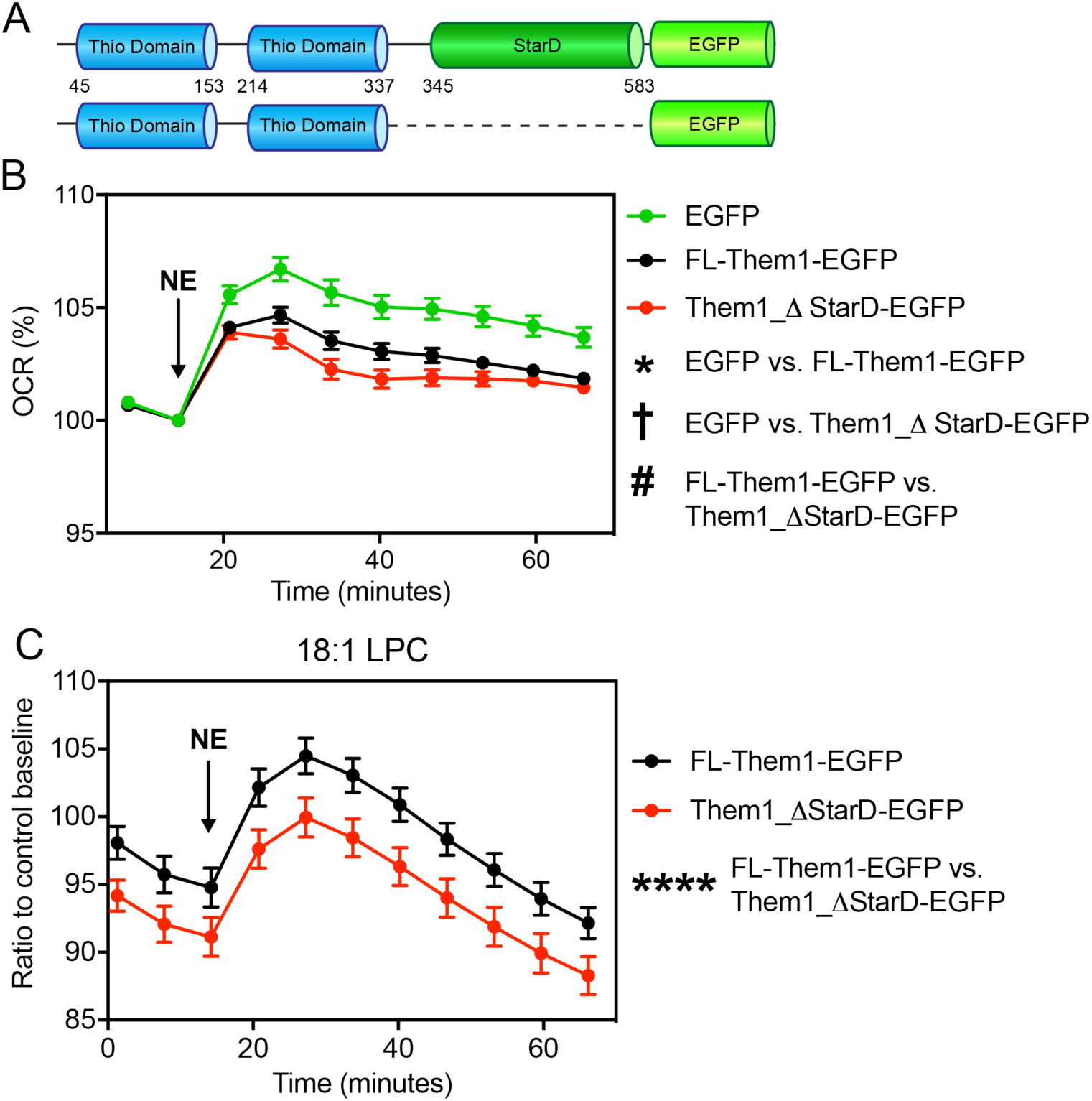
LPC 18:1 inhibits Them1 mediated suppression of thermogenesis in brown adipocytes. *A.* Schematic of adenovirus constructs of full-length Them1 (residues 1-594, top) and Them1_ΔStarD (residues 1-344, bottom) with C-terminal EGFP tags. *B.* OCR of iBAs following stimulation with 1 μM norepinephrine (NE). The iBAs were transduced with Ad-FL-Them1-EGFP (black), Ad-Them1_ΔStarD-EGFP (red), and Ad-EGFP (green). OCR values were normalized by the number of live cell nuclei and displayed as a percentage relative to the basal OCR. Graph shows combined data from 3 independent experiments. Statistical analyses were conducted via 2-way ANOVA with Tukey’s correction. *P<0.001, between EGFP versus FL-Them1-EGFP, †P<0.001, between EGFP versus Them1_ΔStarD-EGFP, # P<0.001, between FL-Them1-EGFP versus Them1_ΔStarD-EGFP. *C.* OCR of iBAs transduced with Ad-FL-Them1-EGFP (black) or Ad-Them1_ΔStarD-EGFP (red) following stimulation with 1 μM NE. Cells were incubated with 25 μM LPC 18:1 or control buffer for one hour prior to the start of experiment. OCR values were normalized by the number of live cell nuclei and displayed as ratios relative to the control baseline OCR for each genetic background. Graphs show combined data from 3 independent experiments. Statistical analyses were conducted via 2-way ANOVA with Tukey’s correction. ****P<0.0001.

It was recently reported that LPC levels, specifically 16:0 and 18:0, were elevated in brown adipocytes upon induction of thermogenesis, and that 16:0 LPC enhanced UCP1 mediated respiration ^40^. Since we identified that 18:1 LPC inhibits Them1 through the StarD (Fig. 6A-B), we hypothesized that 18:1 LPC regulates thermogenesis in brown adipocytes through interaction with Them1. To test this, we incubated iBAs transduced with FL-Them1-EGFP or Them1_ΔStarD-EGFP with 25 μM 18:1 LPC for one hour prior to measuring NE induced respiration. In line with 18:1 LPC inhibition of Them1 through the StarD, respiration of iBAs transduced with FL-Them1-EGFP was enhanced in the presence of 18:1 LPC relative to Them1_ΔStarD-EGFP (Fig. 7C).

### Them1 StarD is necessary for localization to the lipid droplet

To elucidate whether the StarD contributes to Them1 function by altering cellular localization, we visualized Them1 in iBAs stably transduced with the same viral Them1 constructs as above (FL-Them1-EGFP and Them1_ΔStarD-EGFP). FL-Them1-EGFP was primarily localized in puncta near the lipid droplet surface (Fig. 8A-C), as was previously shown by Li et al (REF). Removal of the StarD disrupted lipid droplet localization and led to puncta dispersed throughout the cytosol (Fig. 8D-F). Them1 suppresses thermogenesis in this condensed form, but phosphorylation of the N-terminus disperses Them1 within the cell, which abrogates Them1 mediated inhibition of thermogenesis ^51^. To test whether the StarD alters this phosphorylation-mediated dissolution of Them1, we treated cells expressing just the thioesterase domains (Them1_ΔStarD-EGFP) with phorbol 12-myristate 13-acetate (PMA), a PKC activator that leads to Them1 phosphorylation ^51^. After a 4 h treatment with PMA, Them1 was diffuse, demonstrating that the StarD is not essential for this process (Fig. 8G-H). These data reveal a spatiotemporal role for the Them1 StarD, whereby the StarD is necessary for positioning Them1 puncta near the lipid droplet. However, this process is distinct from the phosphorylation-regulated dynamics between puncta and diffuse Them1 that is critical for the suppression of thermogenesis.

**Figure 8.**
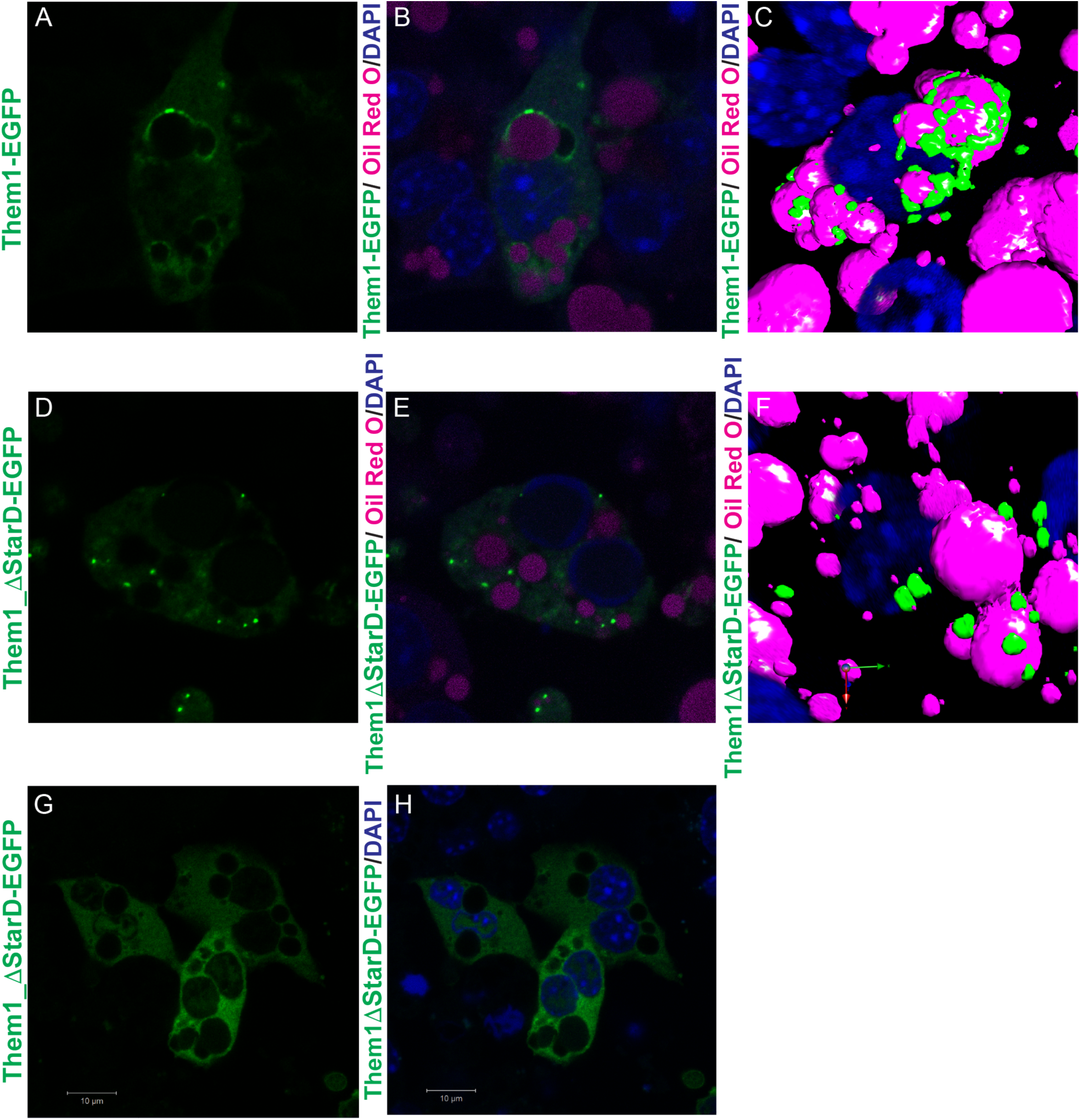
Them1 StarD drives localization to the lipid droplet and is not necessary for Them1 mediated suppression of thermogenesis. *A-F.* Confocal fluorescence microscopy of iBAs cells reconstituted with Ad-FL-Them1-EGFP *(A-C)* and Ad-Them1_ΔStarD-EGFP *(D-F).* Lipd droplets and nuceli were visualized through staining with Oil Red O and DAPI respectively. FL-Them1-EGFP localized with lipid droplets, while Them1_ΔStarD-EGFP did not associate with lipid droplets. *C, F.* 3D rendering of confocal fluorescence microscopy images. G*-H.* PMA treatment of iBAs induced dispression of Them1_ΔStarD-EGFP puncta.

## Discussion

Them1 suppresses thermogenesis in BAT, limiting its capacity to oxidize endogenous fatty acids ^6, 8^. Whereas we previously demonstrated that the C-terminal StarD is necessary for full catalytic activity of the enzyme ^7^, the current study elucidates the multifunctional role of the StarD to act as a lipid sensor to allosterically regulate Them1 activity and spatially localize Them1 near the lipid droplet.

We identified that both long-chain fatty acids and 18:1 LPC bind to the StarD and inversely alter Them1 stability and activity, establishing the StarD as a sensor that has evolved to bind specific lipids to tune enzymatic activity. The allosteric enhancement of activity by myristic and palmitic acid through a feedforward mechanism could drive Them1’s preference to hydrolyze myristoyl-CoA and palmitoyl-CoA ^7^, distinguishing it from other thioesterases. When taken together with phosphorylation-dependent cellular dispersion of Them1 ^51^, the observation that 18:1 LPC as an allosteric inhibitor suggests multiple mechanisms have evolved to suppress Them1 activity in order to enhance thermogenesis. One study showed that induction of thermogenesis in brown adipocytes increased levels of 16:0 LPC, but decreased levels of 18:1 LPC ^40^. However, another study showed that 18:1 LPC levels were increased in browning white adipose tissue of mice treated with a β3-adrenergic agonist ^52^. Them1 potentially senses the nutritional state of these cells through the StarD and regulates its activity to conserve or oxidize lipids. Additionally, 18:1 LPC may spatiotemporally regulate Them1 activity to control lipid droplet membrane development. Since Them1 is localized to the lipid droplet surface, it could interfere with lipid droplet monolayer formation by hydrolyzing fatty acyl-CoAs, which are the substrates used by LPCAT2 to generate phosphatidylcholines for lipid droplet expansion ^39^.

However, this could be prevented through inhibition by LPC, which would be localized at the lipid droplet and is utilized by LPCAT2 to produce phosphatidylcholines.

Our data suggest the mechanism by which fatty acids enhance and 18:1 LPC inhibits Them1 activity is by differential stabilization of the StarD. The C-terminal α-helix of StarD is a gate for ligand binding, remaining unfolded in the apo state, and folding and encapsulating the pocket once ligand binds ^49, 50^. Our molecular dynamic simulations showed myristic acid stabilized this helix, which is apposed to the thioesterase domains. We expect 18:1 LPC destabilizes this helix, which would then destabilize the thioesterase domains. Phospholipids are imperfect pharmacological tools due to their poor pharmacokinetics; however, these findings should aid in the development of small molecule allosteric inhibitors that enhance metabolism. Screening for compounds that destabilize the StarD and in turn inhibit Them1 activity could yield Them1-selective pharmacological tools to treat obesity and related metabolic disorders.

Our confocal fluorescence studies showed the StarD was responsible for localizing Them1 near the lipid droplet in brown adipocytes at a basal state. The StarD of Them1 could directly interact with the lipid droplet membrane or engage in protein-protein interactions at the lipid droplet surface. Previously it was shown that Them1 associated with phosphatidyl inositol-4-phosphate (PIP) through the StarD in a protein-lipid overlay assay, suggesting the StarD of Them1 is capable of directly interacting with a membrane surface ^53^. There were no PIP species that copurified with the StarD in our affinity purification–MS technique, which would only detect high affinity ligands that remain bound through several washing steps; therefore, the StarD may engage in a low affinity interaction with PIP. Recently, it was shown that PIPs are present on the lipid droplet surface, which could potentially explain Them1 localization ^54^. In our fitted model of Them1 in the low-resolution negative stain map, the StarDs are positioned on the exterior of the trimeric complex, where the StarDs could cooperatively bind to the membrane surface, anchoring Them1 puncta to the lipid droplet. Although this localization was shown to be dispensable for Them1-mediated suppression of thermogenesis, the StarD of Them1 could perform other functions at the lipid droplet. For instance, many StarDs are involved in transporting specific lipids to cellular compartments; however, these possibilities remain to be explored ^15, 17, 21^.

In order to properly traffic acyl-CoAs into specific pathways to drive metabolism, the localization and activity of multiple acyl-CoA thioesterases and synthetases must be controlled^55^. The StarD of Them1 allows for fine tuning of Them1 function; regulating both activity and localization. This control is necessary for correct metabolism in BAT, enabling thermogenesis or preserving resources when needed.

## Experimental Procedure

### Materials and reagents

Chemicals were purchased from Sigma-Aldrich, Polysciences Inc., Cayman Chemical, and Avanti Polar Lipids. Cell culture media was purchased from Gibco. The vector for His-tagged tobacco etch virus (TEV) was a gift from John Tesmer (University of Texas at Austin). The pMCSG7 (LIC_HIS) vector was provided by Dr. John Sondek (University of North Carolina at Chapel Hill). The hSTARD14 pNIC28-Bsa4 vector was provided by Dr.

Nicola Burgess-Brown (Structure Genomics Consortium). The pLVX-IRES-ZsGreen1 vector for stable cell line development was donated by Dr. Rafi Ahmed (Emory University). DNA oligonucleotide primers were synthesized by IDT (Coralville, IA).

### Cell Culture

The *HEK293T* cells, which were used to generate the lentivirus, were purchased from the American Type Culture Collection and grown in Dulbecco’s Modified Eagle Medium supplemented with 10 % fetal bovine serum (Atlanta Biologicals) and 1% penicillin– streptomycin. Cells were maintained using standard culture conditions. The Freestyle *HEK293F* cells, which were stably transduced, were purchased from Gibco and grown in Freestyle 293 expression medium supplemented with 1x antibiotic-antimycotic (Gibco). Freestyle *HEK293F* cells were grown in suspension culture using glass flasks and a Benchmark Scientific Orbi-Shaker CO2 shaking at 120 rpm. Cells were maintained at a density of 0.1 – 4 million cells/ milliliter using standard culture conditions. The immortalized brown adipocytes (iBAs) used for localization and Seahorse experiments were a gift from Dr. Bruce Spiegelman (Harvard University). Prior to differentiation, iBAs were grown in Dulbecco’s Modified Eagle Medium supplemented with 20 % fetal bovine serum and 1% penicillin–streptomycin. Differentiation of iBAs was induced through incubation in Dulbecco’s Modified Eagle Medium supplemented with 10 % fetal bovine serum, 1% penicillin–streptomycin, 20 nM insulin, 1 nM triiodo-L-thyronine (T3), 1 μM rosiglitazone, 2 μg/ml dexamethasone, 125 μM indomethacin, and 500 μM 3-Isobutyl-1-methylxanthine (IBMX). After 48 hours, differentiated iBAs were transferred to maintenance medium containing Dulbecco’s Modified Eagle Medium supplemented with 10 % fetal bovine serum, 1% penicillin–streptomycin, 20 nM insulin, 1 nM triiodo-L-thyronine (T3), 1 μM rosiglitazone, and 1 μM norepinephrine (NE). Cells were ready for experimentation after 48 hours in maintenance medium.

### Protein expression and purification

The *Homo sapiens* Them1 START domain (residues 339 – 594 of isoform 2) in the pNIC28-Bsa4 vector was transformed in *Escherichia coli* strain BL21 (DE3) cells that were additionally transformed with the pG-Tf2 vector (codes groES-gorEL-tig chaperones). The START domains were expressed as a His_6_ fusion containing a tobacco etch virus protease cleavage site to facilitate tag removal. Cultures (1 liters in TB) were grown to an *A_600_* of ∼0.6 -0.8 and chaperone transcription was induced with 5 ng/mL tetracycline HCl at 18 °C for one hour, followed by START domain induction with 0.5 mM isopropyl β-d-1-thiogalactopyranoside at 18 °C for ∼18 hours. Cell mass was harvested, lysed through sonication in a buffer containing 20 mM Tris HCl pH 7.4, 500 mM NaCl, 25 mM imidazole, 5% glycerol, lysozyme, Dnase A, 0.1 % Triton X-100, 5 mM beta-mercaptoethanol, and 100 uM phenylmethylsulfonyl fluoride. The START domain was purified by nickel affinity chromatography and the His tag was cleaved by tobacco etch virus protease at 4 °C overnight with simultaneous dialysis into a buffer containing 20 mM Tris HCl pH 7.4, 500 mM NaCl, and 5% glycerol when necessary. Cleaved START domain was purified from His tag through nickel affinity chromatography followed by size exclusion chromatography (SEC) using a HiLoad 16/60 Superdex 75 column.

The *Mus musculus* Them1 thioesterase domains (ΔNterm-Them1_ΔStarD) (residues 43 – 365) in the pMCSG7 vector were transformed into *Escherichia coli* strain BL21 (DE3) pLysS cells. The thioesterase domains were expressed as a His_6_ fusion containing a tobacco etch virus protease cleavage site. Cultures (1 liters in LB) were grown to an *A_600_* of ∼0.6 - 0.8, and thioesterase domain expression was induced with 0.5 mM isopropyl β-d-1-thiogalactopyranoside at 18 °C for ∼18 hours. Cell mass was harvested, lysed through sonication in a buffer containing 20 mM Tris HCl pH 7.4, 500 mM NaCl, 25 mM imidazole, 5% glycerol, lysozyme, Dnase A, 5 mM beta-mercaptoethanol, and 100 uM phenylmethylsulfonyl fluoride. The thioesterase domains were purified by nickel affinity chromatography followed by SEC using a HiLoad 16/60 Superdex 200 column.

Wild-type *Mus musculus* Them1 containing both thioesterase domains and START domain (ΔNterm-Them1) (residues 43 – 594) was cloned along with a N-terminal His_6_ tag followed by a tobacco etch virus protease cleavage site into the pLVX-IRES-ZsGreen1 lentiviral vector. Polyethylenimine, linear (MW 25,000) (Polysciences Inc.), was used to transfect the Them1 pLVX-IRES-ZsGreen1 vector along with the lentiviral packaging (Pax2) and envelope (MD2G) vectors at a mass ratio of 4:2:1 respectively into *HEK293T* cells according to manufacturer’s instructions. After 48 and 72 hours, culture supernatant was collected and viral particles were precipitated through incubation with 10 % PEG 8000, 0.3 M NaCl, and PBS at 4 °C overnight. Viral particles were harvested through centrifugation at 1,500 x g for 30 minutes, decanted, and resuspended in DMEM. Several serial dilutions of lentivirus in DMEM were used to transduce Freestyle *HEK293F* cells. After 72 hours, multiplicity of infection (MOI) was determined for cell lines through measuring GFP expression in limiting dilutions using a flow cytometer. Them1 (MOI of 50) grown in suspension culture was harvested at a cell density of 2 million cells/ milliliter. Cells were lysed through sonication in a buffer containing 20 mM Tris HCl pH 7.4, 500 mM NaCl, 25 mM imidazole, 5% glycerol, lysozyme, Dnase A, 0.1 % Triton X-100, 5 mM beta-mercaptoethanol, and 100 uM phenylmethylsulfonyl fluoride. Them1 was purified by nickel affinity chromatography followed by SEC using a HiLoad 16/60 Superdex 200 column.

### Analytical Ultracentrifugation

Analytical ultracentrifugation experiments were carried out using a Beckman Coulter ProteomeLab™ XLI analytical ultracentrifuge equipped with both absorbance and interference optics and a four-hole An-60 Ti analytical rotor. Sedimentation velocity experiments were carried out at 10 °C and 50,000 rpm (200,000 × g) using 120-mm two-sector charcoal-filled Epon centerpieces with quartz windows. Each sample was scanned at 0-min time intervals for ∼ 200 scans. ΔNterm-Them1 was run at ∼0.5 mg/mL in buffer containing 20 mM bis-Tris pH 8.5, 500 mM NaCl, and 0.5 mM tris(2-carboxyethyl)phosphine (TCEP). Sedimentation boundaries were analyzed by the continuous distribution (c(s)) method using the program SEDFIT ^56^. The program SEDNTERP, version 1.09, was used to correct the experimental s value (s*) to standard conditions at 20 °C in water (s_20,w_) and to calculate protein partial specific volume ^57^. Corrected s_20,w_ was used for molecular weight calculation.

### Lipid exchange with affinity-mass spectrometry

Bovine liver lipid extracts (Avanti Polar Lipids) suspended in chloroform was dried with nitrogen gas, followed by drying with a vacuum desiccator for at least an hour. Liposomes were generated through resuspending dried lipids in 20 mM Tris pH 7.4, 150 mM NaCl, and 5 % glycerol, agitating for one hour at room temperature, and sonicating for one hour in a bath sonicator. Purified His-tagged StarD and His-tagged maltose binding protein were incubated with liver lipid vesicles at 4 °C overnight with total lipid concentration in five-time excess to protein. Non-specifically bound lipids were removed through further purifying proteins with nickel affinity chromatography and SEC using a HiLoad 16/60 Superdex 75 column. Prior to running SEC, the column was cleaned by thoroughly washing with 70 % ethanol, followed by washing with deionized water and equilibration with 20 mM Tris pH 7.4, 150 mM NaCl, and 5 % glycerol. Glass washed three times with chloroform was used to collect, handle, and store samples proceeding SEC. Equal mass samples of StarD and maltose binding protein were collected in triplicate ranging from 0.5 – 1.0 mg depending on the experiment. SEC buffer was added to samples to make volume equal for all samples.

Additionally, three SEC buffer samples of equal volume were collected as a negative control. Lipids were extracted from samples using the Bligh and Dyer method ^58^. Samples were solubilized in 400 μL 1:1 v/v methanol: chloroform mix and spiked with either deuterated palmitic acid-d2 (Cayman Chemical) or deuterated lipid standards (SPLASH II LIPIDOMIX, Avanti Polar, Alabaster, Alabama). Fatty acids were analyzed using direct infusion mass spectrometry in Enhanced MS (EMS) mode over one minute and averaged. The profile mode data was collected in negative mode at scan rate of 10000 Da/s within the mass range 100-1000. The instrumental parameters used were as follows: curtain gas-20, CAD-Low, Ion spray voltage--4500, temperature – 350 °C and declustering potential --100. The data was processed by extracting the peak area from the mass spectrum data and lipid was identified using mass with M-H adduct. Fatty acids were normalized to deuterated palmitic acid standard to quantify fatty acid species. The data is graphically represented as an average of three technical replicates +/-SEM. Phospholipids were analyzed through injecting ten µL of each sample into the mass spectrometer for flow injection analysis. Several precursor ion scan methods were applied to specifically target phosphatidylcholine, phosphatidylethanolamine, sphingomyelin, ceramide, phosphatidylserine and phosphatidylinositol in the samples. Mass spectrometry data was collected for each precursor ion scan over one minute and averaged. The data was analyzed utilizing LipidView 1.2 (SCIEX, Framingham, MA, USA) with the following processing parameters: polarity-positive, precursor ion scan and neutral loss scan, mass tolerance-0.5, minimum S/N-10, minimum % intensity-0. With these settings, the data were smoothed, deisotoped and searched for the peak list within a *m/z* range of 100-1000 and chromatographic range of 0.4-1.2 minute. The peak list generated through LipidView 1.2 was further interrogated using the LIPIDMAPS online tool. The identified lipids were semi quantified using deuterated lipid standards. Zero concentrations are represented as not determined (ND). The data is graphically represented as an average of three technical replicates +/-SEM. Statistical analyses were conducted using 2-way ANOVA with Sidak’s multiple comparisons test.

### Microscale thermophoresis

His-tagged *Homo sapiens* Them1 START domain was labeled using the Monolith His-Tag Labeling Kit RED-tris-NTA (NanoTemper Technologies). The labeling reaction was performed according to the manufacturer’s instructions in PBS supplemented with 0.5 % Tween 20 at a concentration of 100 nM protein (molar dye:protein ratio = 1:1) at room temperature for 30 min. Fatty acids were dissolved in ethanol at 10 – 20 mM, and diluted in PBS supplemented with 0.5 % Tween 20 in a series of 16 1:1 dilutions, producing ligand concentrations ranging from 12.2 nM – 800 μM, with a constant final ethanol concentration of 4 %. Each ligand dilution was mixed with one volume of labeled START domain, which led to a final concentration of START domain of 50 nM and final ligand concentrations ranging from 6.1 nM to 400 μM. START domain was incubated with fatty acid overnight at 4 °C, then loaded in standard Monolith NT.115 Capillaries (NanoTemper Technologies). MST was measured using a Monolith NT.115 instrument (NanoTemper Technologies) at an ambient temperature of 25°C. Instrument parameters were adjusted to 40 % LED power and medium MST power. Data of three independently pipetted measurements were fitted with a non-linear regression model in GraphPad Prism 8.0 using the signal from an MST-on time of 5 s.

### Crystallization, data collection, structural refinement

Pure *Homo sapiens* Them1 START domain was incubated with myristic acid in 10-fold excess and concentrated to 10 mg mL^-1^ in 30 mM HEPES pH 7.5, 300 mM NaCl, 10 % glycerol, 0.5 mM tris(2-carboxyethyl)phosphine (TCEP). Crystals of START domain were grown over two weeks via hanging drop vapor diffusion at 4 °C from solutions containing 1 μL START domain and 1 μL mother liquor (0.1 M Tris HCl pH 9.4 and 27 % PEG 8000). Crystals were cryoprotected by immersion in 0.1 M Tris HCl pH 9.4, 27 % PEG 8000, and 20 % glycerol and flash frozen with liquid nitrogen. Data were collected remotely from the Southeast Regional Collaborative Access Team at the Advanced Photon Source, 22ID beamline (Argonne National Laboratories, Chicago, IL). Data were processed and scaled using HKL-2000 (HKL Research, Inc., Charlottesville, VA) ^59^ and phased by molecular replacement using Phaser-MR (Phenix, Berkeley, CA) ^60^. The structure was phased using a previously solved crystal structure of Them1 START domain (3FO5) as a search model ^32, 61^. Structure refinement and validation was performed using PHENIX (Phenix, Berkeley, CA) (version 1.11.1), and model building was performed in COOT (MRC Laboratory of Molecular Biology, Cambridge, UK) ^60, 62^. PyMOL (version 1.8.2; Schrödinger, New York, NY) was used to visualize structures and generate figures. Structure is deposited in PDB with ID: 6VVQ.

### Acyl-CoA thioesterase activity assay

Myristic acid, palmitic acid, stearic acid, palmitoleic acid, and 18:1 lysophosphatidylcholine were dissolved in ethanol to a concentration of 1.78 mM (3.55 mM for 18:1 LPC at final 50 μM). Purified *Mus musculus Δ*Nterm-Them1 or thioesterase domains at a concentration of 1.42 μM were incubated with 429 μM 5,5′-dithiobis-(2-nitrobenzoic acid) (DTNB) and 35 μM fatty acid or 18:1 lysophosphatidylcholine (71 μM for 18:1 LPC at final 50 μM) for 30 minutes at 37 °C in assay buffer containing 30 mM Hepes pH 7.5, 150 mM NaCl, and 5 % glycerol; ethanol concentration was 2 % across samples. Myristoyl-CoA (Sigma) was dissolved in 10 mM MES pH 5.5 to a concentration of 5 mM and further diluted in assay buffer. Myristoyl-CoA was added to protein-DTNB-lipid mixture to initiate reaction in a total reaction volume of 200 μL/ well, with final protein concentration at 1 μM, DTNB at 300 μM, lipid at 25 μM or 50 μM, and myristoyl-CoA ranging 0 – 20 μM. Plates were immediately introduced into a 37 °C temperature-controlled Synergy Neo 2.0 (BioTek) plate reader. Absorbance readings at 412 nm were read every 10 seconds for 1 hour. Enzyme initial velocities (*V_0_*) were calculated through fitting a line to the rise in product formation in the early time points using GraphPad Prism 8.0 for each substrate concentration. The initial velocities were plotted against substrate concentrations and fitted with the Michaelis–Menten equation to yield the maximum velocity (V_max_) and Km (the Michaelis constant) using GraphPad Prism 8.0. Values of kcat were calculated as kcat = Vmax/[E]. Each experiment was conducted with two technical replicates for each sample and repeated three times.

### Negative Stain Electron Microscopy

Freshly purified *Mus musculus Δ*Nterm-Them1 (residues 43 – 365) after SEC into a buffer containing 30 mM Hepes pH 7.5, 300 mM NaCl, and 0.5 mM TCEP was used for negative staining. Briefly, 4 ul of ΔNterm-Them1 was adsorbed on a carbon coated Cu–400 mesh grid (TedPella) grid for 1 minute and excess liquid was blotted with Whatman Filter paper 4, washed twice with 20 ul drop of water and stained with 0.75 % Uranyl formate for 30 sec, blotted and air dry. Negatively stained Them1 was imaged on Talos 120 C Microscope operating at 120 kV with Lab6 cathode at pixel size of 1.56 Å. Micrographs were recorded at low dose condition on the Ceta 16M camera (ThermoFisher).

### 2D classification and 3D Reconstruction

32000 particles were auto picked from 267 micrographs using EMAN2 e2boxer. Initial 2D classification was done using ISAC2 program in Sphire Package ^63^. Final 6907 particle stack was imported to Relion 3.0 for further analysis. Ab-inito model reconstruction and 3D refinement was done using C1 and C3 symmetry respectively. Final 4457 particles were selected which gave the resolution of 23 Å with 0.5 FSC criteria. Final 3d volume with C3 symmetry was back projected and compared to 2D classes using EMAN2 for model validation. Model Visualization and analysis was done in Chimera.

### Molecular Dynamics Simulations

A model of Them1 monomer was prepared for molecular dynamics (MD) simulations. Them1 was prepared by aligning the thioesterase domains of ACOT12 bound to ADP and CoA (PDB code: 4MOB) and our myristic acid bound structure of the StarD. Using this model, apo and myristic-acid bound Them1 complexes were created. Complexes were solvated using an octahedral box of TIP3P water ^64^ with a 10 Å buffer surrounding the complexes. Complexes were first neutralized and then adjusted to a final concentration of 150 mM NaCl by the addition of Na+ and Cl-ions. All complexes were prepared using xleap in AmberTools ^65^ and the parm99 forcefield ^66^ in Amber14 ^67^. Parameters for ADP, CoA and myristic acid were obtained using Antechamber ^68^ in AmberTools. Using 5000 steps of steepest descent followed by 5000 steps of conjugate gradient, systems were minimized in two rounds. In the first round, restraints of 500 kcal/mol-A^2^ were applied to all solute atoms. In the second round, restraints were removed from protein atoms and only maintained for the ligands. Systems were then heated from 0 to 300 K using a 100-ps run with constant volume periodic boundaries and restraints of 10-kcal/mol.A^2^ applied to ligands (i.e. myristic acid, CoA and ADP). Equilibration was performed using 12 ns of MD in the NPT ensemble with 10-kcal/molA^2^ restraints on small molecule atoms. Restraints were reduced to 1- kcal/molA^2^ and equilibration performed for an additional 12 ns. All restraints were removed and 500-ns production simulations performed for each system. All bonds between heavy atoms and hydrogens were fixed with the SHAKE algorithm ^69^, allowing the use of a 2-fs time step. Long-range electrostatics were evaluated with a cutoff of 10 A.

For analysis, 25000 evenly-spaced frames were obtained from each simulation. The CPPTRAJ ^70^ module of AmberTools was used for structural averaging and calculations of root mean square fluctuations (RMSFs). Dynamic networks were produced for each system using the NetworkView plugin ^71^ of VMD ^72^. Networks are constructed by defining all protein C-α atoms as nodes, using Cartesian covariance to measure communication within the network. Pairs of nodes that reside within a 4.5-Å cutoff for >75% of the simulation are connected via an edge.

Edge weights are inversely proportional to the covariance between the nodes. Networks were subsequently partitioned into communities using the Girvan-Newman algorithm ^73^. Communities represent a group of nodes undergoing correlated motions. The minimum number of communities possible was generated while maintaining at least 98% maximum modularity.

### Differential scanning fluorimetry (DSF)

Myristic acid, palmitic acid, stearic acid, and 18:1 lysophosphatidylcholine were dissolved in ethanol to a concentration of 2.5 mM. Purified His-tagged *Homo sapiens* Them1 START domain and His-tagged *Mus musculus* Them1 thioesterase domains at a concentration of 10 μM were incubated with 50 μM fatty acid or 18:1 lysophosphatidylcholine for 30 minutes at room temperature; ethanol concentration was constant at 2 % across samples. SYPRO orange dye (Invitrogen) was then added at a 1:1000 dilution. Reactions were heated at a rate of 0.5 °C per minute, using a StepOne Plus Real Time PCR System (ThermoFisher). Fluorescence was recorded at every degree using the ROX filter (602 nm). Technical triplicates were analyzed by first subtracting baseline fluorescence (ligands + SYPRO with no protein) and then fitting the curves using the Bolzman equation (GraphPad Prism 8.0) to determine the Tm. Experiment was performed with three replicates and the Tm’s of different ligands were analyzed in Prism 8.0 with one-way ANOVA and Dunnett’s multiple comparisons test.

### Hydrogen-deuterium exchange mass spectrometry

HDX-MS experiments were performed on Them1 and the thioesterase domains in two replicates using a Waters nanoACQUITY UPLC HDX system coupled with a Synapt G2-Si mass spectrometer (Waters Corp, Milford, MA). The samples of ΔNterm-Them1 and ΔNterm-Them1_ΔStarD were prepared in 30 mM Hepes pH 7.5, 150 mM NaCl, and 0.5 mM TCEP at a final protein concentration of 6 μM. A PAL system autosampler (LEAP Technologies, Carrboro, NC) mixed the protein samples 1:7 (v/v) with 99.9% D2O-containing buffer (10mM phosphate buffer, pD 7.0) at 20°C for variable time points between 0 and 14,580 seconds before quenching the reaction with an equal volume of pre-chilled quenching buffer (100 mM Na2PO4 pH 2.5, 5% Formic Acid, and 2% Acetonitrile) at 1°C. The quenched samples were digested with a Waters Enzymate BEH Pepsin Column (2.1 X 30 mm). Peptic peptides were then separated using a Waters ACQUITY UPLC BEH C18 column (1.7 µm 1.0 X 100 mm) at a flow rate of 40 µL/min for 11 minutes in a 5-85% linear gradient with a mobile phase of acetonitrile and 0.1% formic acid at 1°C. The mass spectrometer operated with the electrospray ionization source in positive ion mode, and data were acquired in resolution mode. A reference lock-mass of leucine enkephalin (Waters, Milford, MA) was acquired during sample data collection for internal calibration. Peptides were sequenced and identified through database searching of *Mus musculus* Them1 (residues 43 – 594) in ProteinLynx Global Server (ver. 3.0.3).

### Confocal fluorescence microscopy

One day following differentiation, iBAs were transduced with adenovirus containing eGFP, *Mus musculus* full-length Them1-eGFP (residues 1-594), or Them1_ΔStarD-eGFP (residues 1-344) at a multiplicity of infection of 40. Three days following transduction, iBAs cells were washed three times with phosphate buffered saline (PBS) and then fixed with 4% paraformaldehyde in PBS for 20 min at room temperature. Cells were washed twice before staining with 0.3% Oil Red O solution in 60% isopropanol for 2 min at room temperature. The cells were washed twice with PBS and then mounted with ProLong Diamond Antifade Mountant with DAPI (ThermoFisher) to identify nuclei. The localization of fluorescence signal in cultured iBAs cells was evaluated using a Zeiss LSM880 confocal microscope system. The following wavelengths were used to image cellular components: the 405 nm laser was used for DAPI to stain nuclei; the 488 nm laser was used to evaluate EGFP that was linked to Them1; and the 543 nm laser was used to visualize lipid droplets that were stained with Oil Red O. Images were acquired through 2 µm z-stack slices at high resolution and assembled using Volocity image processing software.

### O_2_ consumption rates

OCR values were measured in iBAs using an Seahorse XFe96 analyzer (Seahorse Bioscience; North Billerica, MA, USA). iBAs were plated into Seahorse XF96 cell culture plate (Agilent) at a density of 1,000 cells / well and differentiated as described above.

One day after induction of differentiation, cells were transduced with adenovirus containing eGFP, *Mus musculus* full-length Them1-eGFP (residues 1-594), or Them1_ΔStarD-eGFP (residues 1-344) at a multiplicity of infection of 40. Three days post-transduction, iBAs were incubated in the absence of CO_2_ for 1 h at 37 °C in Krebs–Henseleit buffer (pH 7.4) containing 0.45 g/L glucose, 111 mM NaCl, 4.7 mM KCl, 2 mM MgSO_4_-7H_2_O, 1.2 mM Na_2_HPO_4_, 5 mM HEPES and 0.5 mM carnitine (Sigma–Aldrich). For LPC experiments, 16:0 and 18:1 LPC (Avanti Polar Lipids) were solubilized in DMSO at 10 mM, and further diluted into Krebs Henseleit buffer to a final concentration of 25 μM, allowed to incubate with cells at 37 °C for one hour prior to experimentation. OCR values were measured before and after the exposure of cells to 1 μM NE. OCR was normalized with total live cell count calculated through staining live cells with NucRed Live probe and measuring fluorescence with SpectraMax i3X plate reader at ex/em 625/715 nm. Three independent replicates were analyzed and compared with two-way ANOVA and Tukey’s comparison test in Prism 8.0.

## Acknowledgements

The authors thank Dr. Peter E. Prevelige at UAB for his assistance with HDX-MS, Dr. Pete Lollar and Dr. Anamika Patel at Emory University for their assistance with AUC, and Dr. Bruce Spiegelman of the Dana-Farber Cancer Institute at Harvard Medical School for providing the iBAs cells. This work was supported by the National Institutes of Health (RO1 DK 103046 to D.E.C., S.J.H, and E.O.). M.C.T. was funded by the T32 GM008602 NIH Pharmacology Training Grant. Crystallographic data were collected at Southeast Regional Collaborative Access Team (SER-CAT) 22-ID beamline at the Advanced Photon Source, Argonne National Laboratory, and was supported by the United States Department of Energy, Office of Science, Office of Basic Energy Sciences, under contract W-31–109-Eng-38. This study was supported in part by the Emory Integrated Lipidomics Core (EILC), which is subsidized by the Emory University School of Medicine and is one of the Emory Integrated Core Facilities. Additional support was provided by the Georgia Clinical & Translational Science Alliance of the National Institutes of Health under Award Number UL1TR002378. The content is solely the responsibility of the authors and does not necessarily reflect the official views of the National Institutes of Health.

## Author Contributions

M.C.T. performed affinity purification-MS, ligand binding assays, X-ray crystallography, thioesterase assays, negative stain electron microscopy, DSF assays, and wrote the paper; N.I. did Seahorse experiments; Y.L. performed cellular localization studies; M.K. assisted with affinity purification-MS and data analysis; C.D.O. performed MD simulations; P.J. did negative stain electron microscopy; A.A. assisted with thioesterase assays; S.J.H. assisted with experimental design; D.E.C. assisted with experimental design and edited the paper; E.A.O. mentored M.C.T., assisted with experimental design and data analysis, and edited the paper. All authors reviewed and edited the final manuscript.

## Materials & Correspondence

Correspondence and requests for materials should be addressed to E.A.O.

---

**Supplemental Table 1.**
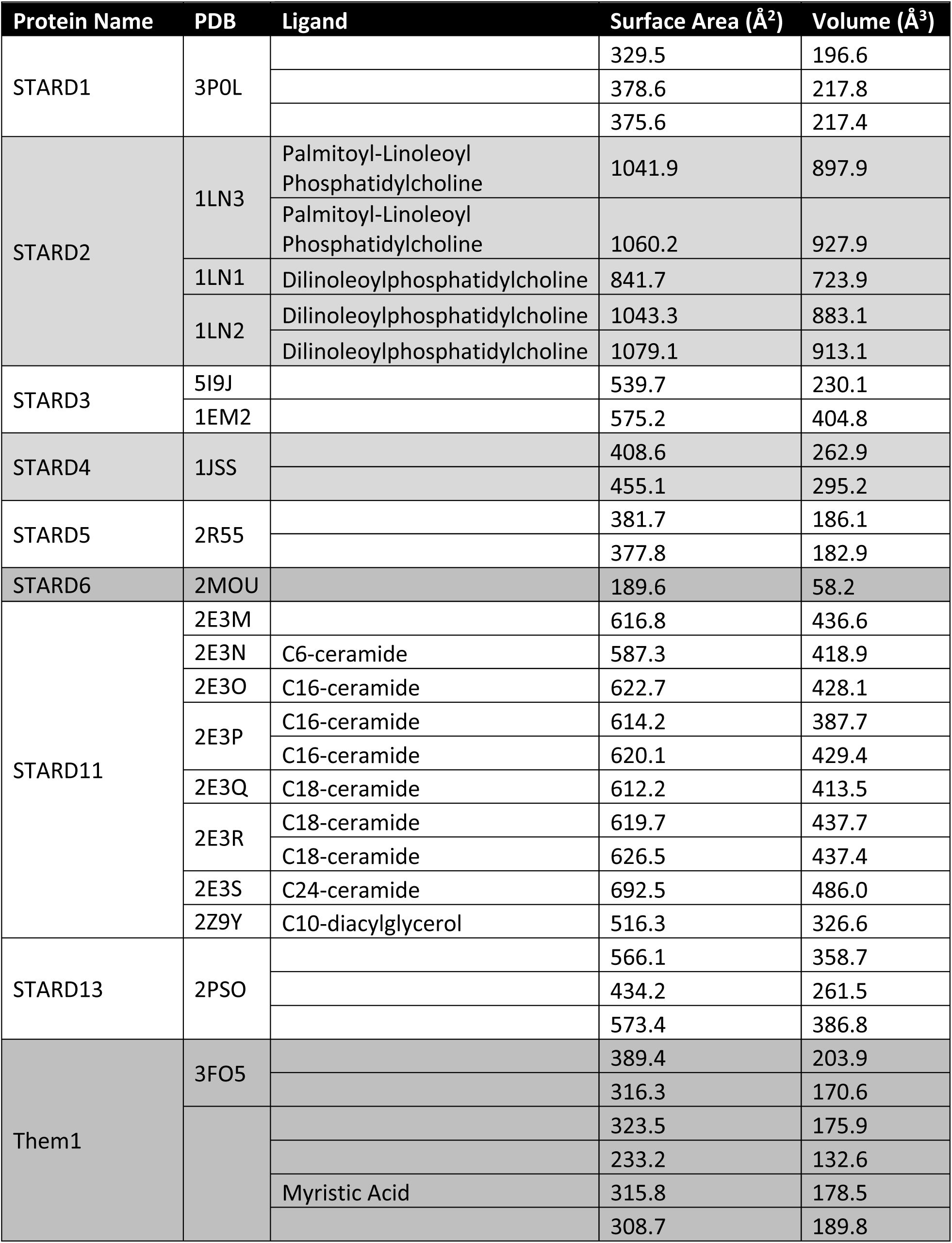

**Supplemental Figure 1.**
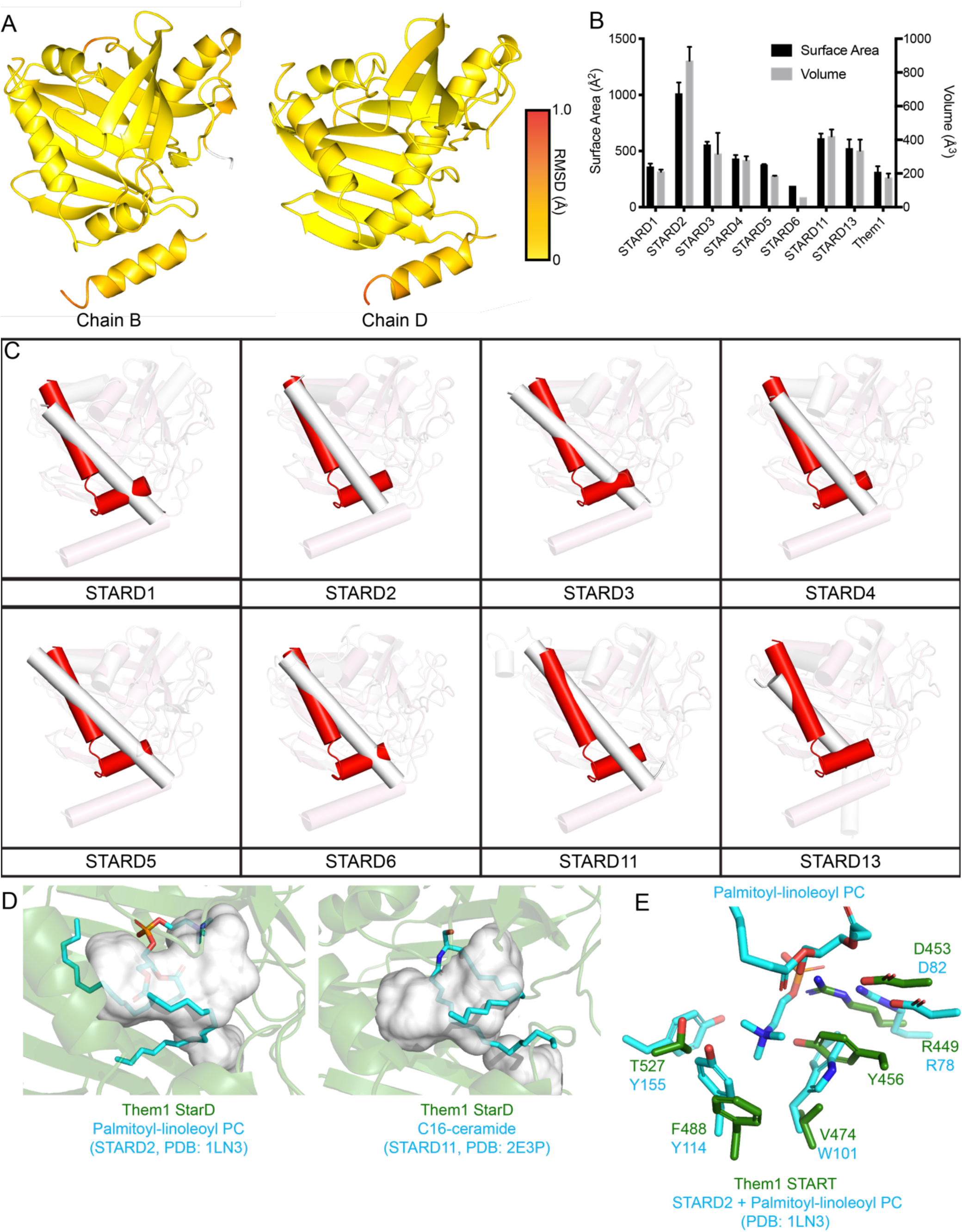
Structure of Them1 StarD suggests small lipids bind. *A.* ProSMART analysis conducted to determine r.m.s.d. between Cα backbone of myristic acid bound StarD and apo-StarD monomers (chain B and D). Root mean square deviations (range: 0–1.0 Å) between monomers were mapped onto chains B or D of the StarD structure with a color scale depicting low (yellow) to high (red) deviations. Unaligned regions are colored in white. *B.* Graphical representation of Table 3. Bars depict average surface area (black) and volume (gray) for each StarD structure as determined from the CASTp server ^74^. Error bars display standard error of the mean. *C*. Alignment of Them1 StarD structure (red) with known structures of all other StarDs (white). The C-terminus of each comparison is emphasized to display structural dissimilarity. *D.* Structural alignment of the StarD of Them1 (green) with StarD2 bound to palmitoyl-linoleoyl phosphatidylcholine (left; PDB code: 1LN3) and StarD11 bound to C16-ceramide (right; PDB code: 2E3P). Only ligands of StarD2 and StarD11 displayed as cyan spheres. Ligands from StarD2 and StarD11 structures do not fit into the interior cavity of the StarD of Them1 that is colored white. *E.* Zoomed in view of the binding pocket of StarD2 (cyan, PDB code: 1LN3) and the StarD of Them1 (green) displaying residues surrounding palmitoyl-linoleoyl phosphatidylcholine from the StarD2 structure. The Them1 StarD contains conserved R449 and D453 that are also found in StarD2, that could participate in electrostatic interactions with the phosphate group of the PC. The StarD of Them1 lacks the aromatic cage present in StarD2 (W101, Y114, Y153), though it contains Y456 that could occupy the same space as W101 in StarD2.

**Supplemental Figure 2.**
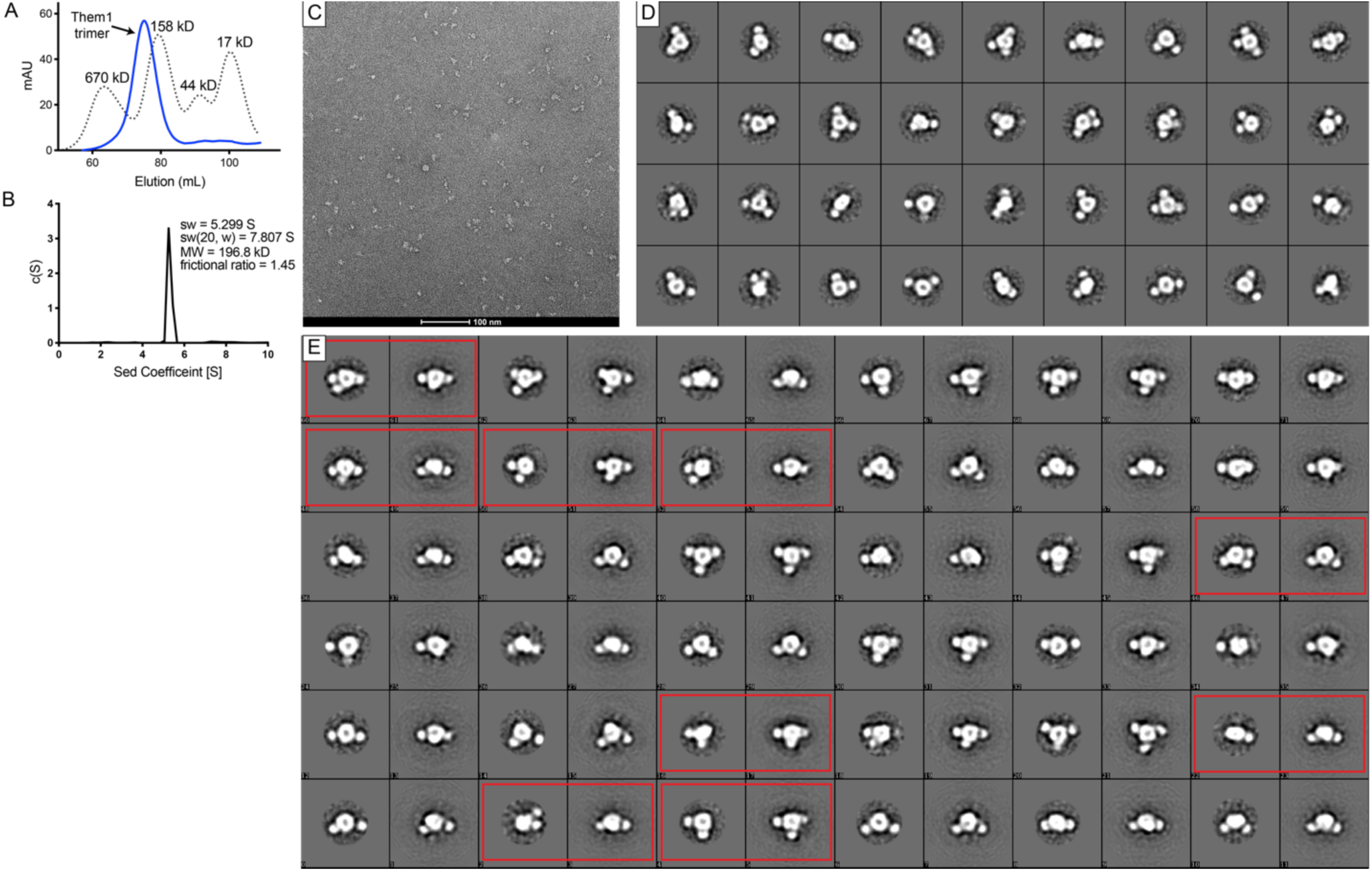
Negative stain single particle electron microscopy of Them1. *A-B*. ΔNterm-Them1 purifies as a homogenous trimeric complex (∼196.5 kD) as determined by size exclusion chromatography *(A)* and analytical ultracentrifugation *(B). A.* Size exclusion chromatography of ΔNterm-Them1 (blue) and standards (dotted line). *B. c(s)* distribution from sedementation velocity analytical ultracentrifugation of ΔNterm-Them1 complex. *C.* One image of ΔNterm-Them1 stained with uranyl formate spread across carbon coated Cu mesh grid collected on Talos 120 C Microscope at a magnification of 96,000X. *D.* 2D Class averages of Them1 using Relion 3.0. *E.* Comparison of 2D class averages (left) and reprojections (right) of 3D model generated with C3 symmetry to match the class average. Red rectangles highlight when reprojections of 3D model do not match class averages.

**Supplemental Figure 3.**
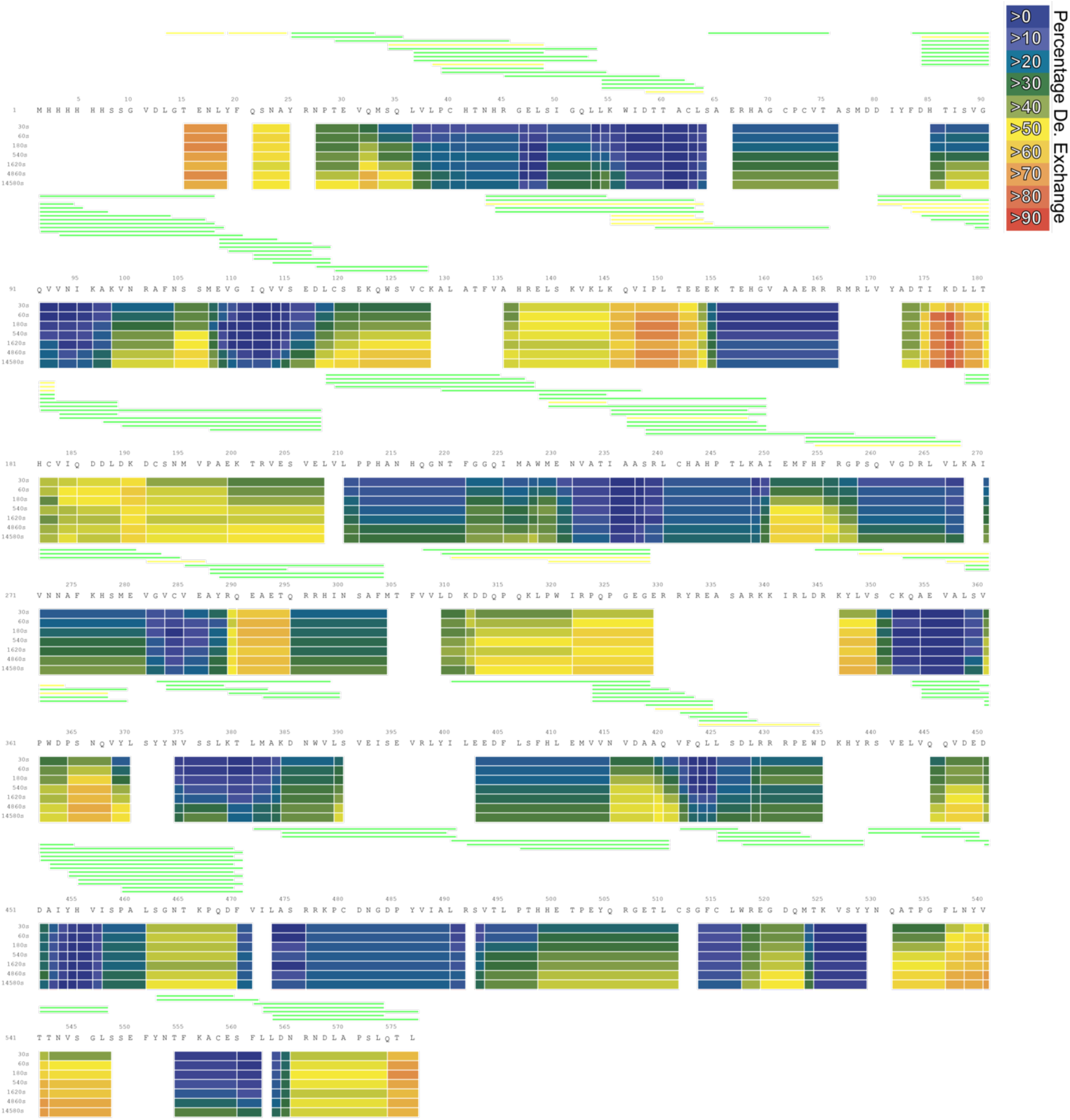
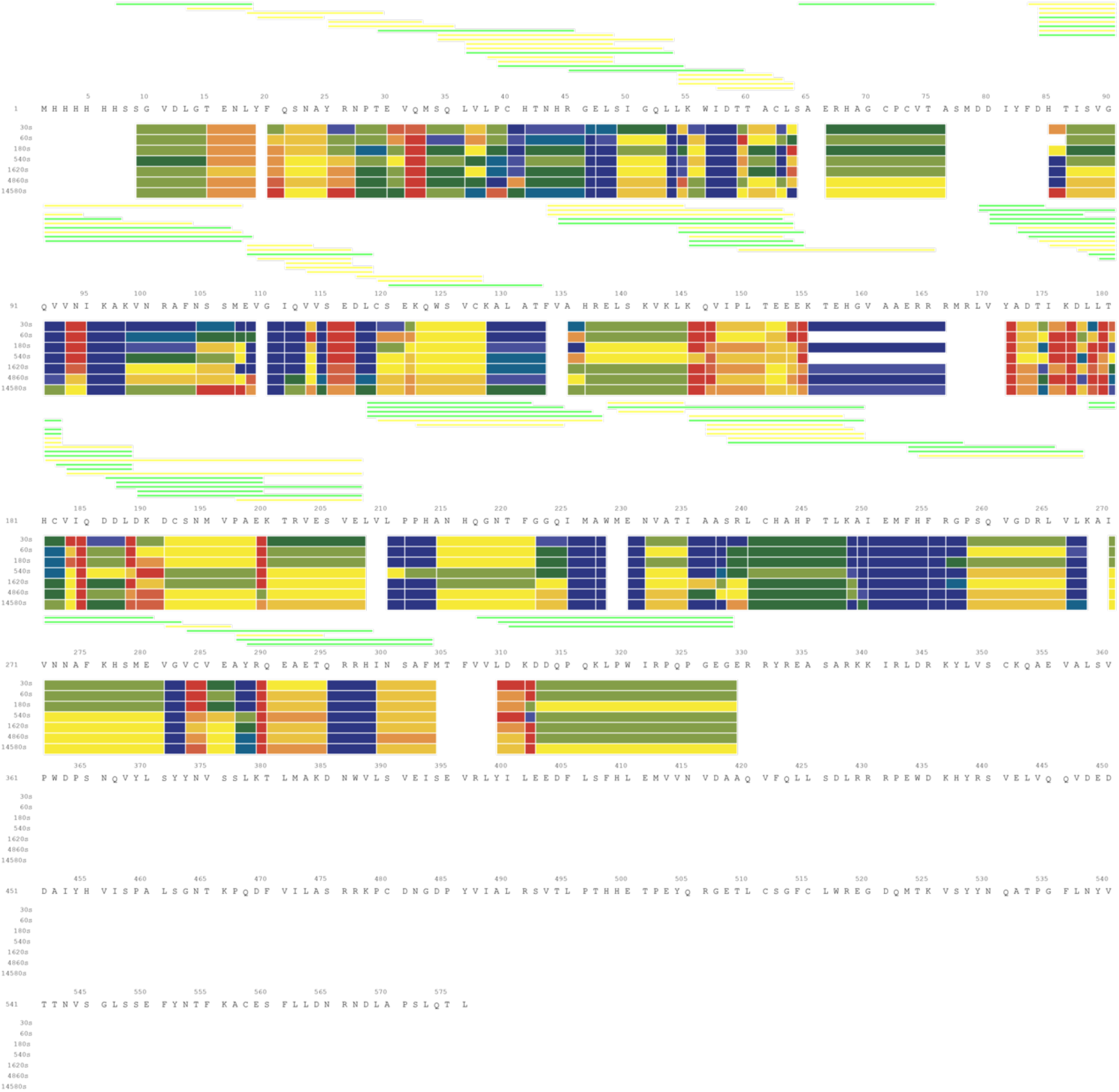
HDX-MS heatmap of Them1 and thioesterase domains. Identified peptides are displayed above sequence and colored according to the identification confidence (green = high confidence; yellow = medium confidence). Below sequence is a heatmap corresponding to the percentage deuterium incorporation across the sequence at each time point. ΔNterm-Them1 (top) is globally more stable than the thioesterse domains alone (bottom).

